# Synthesis, Insertion, and Characterization of SARS-CoV-2 Membrane Protein Within Lipid Bilayers

**DOI:** 10.1101/2023.09.30.560318

**Authors:** Yuanzhong Zhang, Sara Anbir, Joseph McTiernan, Siyu Li, Michael Worcester, Pratyasha Mishra, Michael E. Colvin, Ajay Gopinathan, Umar Mohideen, Roya Zandi, Thomas E. Kuhlman

**Affiliations:** Department of Physics and Astronomy, University of California, Riverside, Riverside, CA USA 92521; Biophysics Program, University of California, Riverside, Riverside, CA USA 92521; Department of Physics, University of California, Merced, Merced, CA USA 95340; Department of Biochemistry, University of California, Riverside, Riverside, CA USA 92521; Department of Chemistry and Biochemistry, University of California, Merced, Merced, CA USA 95340; Microbiology Program, University of California, Riverside, Riverside, CA USA 92521

## Abstract

The membrane protein (M) is the most abundant structural protein in the SARS-CoV-2 virus and functions exclusively as a membrane-embedded homodimer. M protein is required for the formation of the SARS-CoV-2 virus particle and has been shown to interact with the Spike and Envelope proteins, as well as the RNA-packaging Nucleocapsid protein. Our knowledge of M protein is very limited due to its small size and challenges in expressing enough protein for use in structural and biophysical experiments. We report the successful development of a SUMO tag-based expression system to produce and purify significant quantities of M protein, and a method to insert the synthesized dimers into a suspended lipid membrane in a homogeneous orientation. We used AFM and Cryo-EM to image individual membrane-bound M protein dimers and characterize the configurations that they can assume. Our experimental results are in agreement with our molecular dynamics simulations which predict thinning of the membrane around the M protein and a propensity to induce local membrane curvature. Taken together, our results shed new light on M protein properties within the lipid bilayer and suggest mechanisms that could contribute to viral assembly and budding.

## INTRODUCTION

The COVID-19 pandemic has resulted in more than 750 million infections and 6.9 million deaths as of August 2023 ^1^, straining medical and public health systems across the globe. Despite its terrible cost, the COVID-19 pandemic provided the opportunity to demonstrate the effectiveness of combining modern scientific tools, from high-throughput sequencing to computer simulations, to rapidly create treatments and vaccines for a novel infectious agent. However, research on these deadly viruses typically pauses once the immediate danger linked to the diseases they cause diminishes. This, in conjunction with the inherent technical challenges in studying these viruses, some of which are addressed in this paper, has resulted in numerous crucial questions about this dangerous viral family remaining unanswered. In this paper we describe the multidisciplinary application of synergistic tools, including new protein expression and purification methods, advanced spectroscopy, Atomic Force Microscopy (AFM) and Electron Microscopy (EM), as well as coarse-grained and atomistic molecular simulations, to express and characterize the SARS-CoV-2 membrane protein (M Protein) within supported lipid membranes.

SARS-CoV-2 expresses three membrane-bound structural proteins, spike (S), envelope (E), and membrane (M), that play essential roles in the recognition, adhesion, and incorporation of the virus into the host cells, as well as the assembly and budding of newly formed virions. The 222-amino acid M protein is the most common structural protein in SARS-CoV-2 with an estimated 2000 copies in the membrane of each virus particle ^2^. M protein is believed to function exclusively as a membrane-embedded homodimer and is required for the formation of the SARS-CoV-2 virus particle and has been shown to interact with S and E proteins, as well as the RNA-packaging Nucleocapsid (N) protein ^3^.

Compared to the highly studied S protein, relatively little is known about the structure and function of M protein despite its abundance and undisputable crucial role in the budding and formation of the virus ^3, 4^. The limited understanding of M proteins is primarily due to their small size and the difficulties associated with producing adequate quantities for use in structural and biophysical experiments, such as small angle X-ray scattering (SAXS), cryo-EM, etc. Initial structural studies on M protein were based on homology-based predictions of its structure ^5^, but in 2022 two Cryo-EM structures were published. Zhang *et al.* studied homodimers of M protein expressed in human Expi293F cells and embedded in detergent micelles ^3^. To help in the Cryo-EM structure determination, Zhang et al. included antigen binding fragments (Fabs) of mouse monoclonal anti-M protein antibodies ^3^. Their results revealed two distinct M protein structures, a “short” and a “long” form, a result consistent with earlier studies on SARS-CoV M protein using Cryo-EM tomography by Neuman *et al.* ^6^. Dolan *et al.* determined the structure for M protein homodimer of SARS-CoV-2 expressed in Sf9 (insect) cells and embedded in lipid nanodisks ^4^. They found a single M protein structure, similar to Zhang’s short form and suggested that the Fab binding may have mimicked binding to N, S, or other components to stabilize the long form which is otherwise not stable. Both of these studies used atomistic molecular dynamics to establish the stability of their M protein structures in simple lipid membranes.

Despite the important recent progress, many questions remain unanswered about the structure and function of the M protein. These include whether M protein actually exists in two separate stable structures and, if so, whether these have distinct functions, whether the M protein on its own affects the curvature of the virion membrane, or whether that requires binding by other viral proteins, and what are the structure and stoichiometry of M protein’s interactions with the S, E, and N proteins, as well as the viral RNA. Answering these questions will require a collective long-term collaboration involving the entire SARS-CoV-2 scientific community. In this paper, we present our successful creation of a reliable process to express and purify large quantities of M protein and the ability to insert M protein dimers into suspended membranes for biophysical and structural characterization, a capability essential for elucidating the role of M proteins in the formation of SARS-CoV-2.

More specifically we describe our development and optimization of a Small Ubiquitin-related Modifier (SUMO) tag-based expression system to produce and purify significant quantities of M protein. We have demonstrated the purity of our M protein product using western blot and matrix-assisted laser desorption/ionization (MALDI) mass spectrometry and shown that it has low endotoxin activity. We have also developed a method to insert the synthesized M protein dimers into a suspended lipid membrane in a homogeneous orientation. We used Cryo-EM and AFM to image the M protein dimers in the membrane and have performed all-atom molecular dynamics simulations of both the long- and short-form structures of the M protein dimers in a lipid membrane to provide structural information about the protein and the surrounding membrane to aid in interpreting the imaging results. The simulation and microscopy results provide information about the positioning of the M protein dimers in the membrane, its effects on the thickness of the surrounding membrane, and also suggest that M protein dimers alone can potentially induce local curvature in the membrane. Finally, our coarse-grained MD simulations demonstrate that sufficiently strong M-M interaction would be sufficient to drive budding of the SARS-CoV-2 virus.

## RESULTS

### Expression and High Yield Purification of SARS-CoV-2 Membrane (M) Protein in *E. coli*

To assess the level of expression of SARS-CoV-2 Membrane (M) protein in *E. coli*, we constructed a C-terminal fusion of M with the bright blue fluorescent protein mCerulean3. This protein is borne on the high copy number plasmid pUC57 and expressed from the promoter P*lac*T7, inducible with IPTG. Upon transforming this construct into the standard expression strain BL21(DE3), we observed extremely low expression regardless of the degree of induction with IPTG (Fig. 1a).

**Figure 1.**
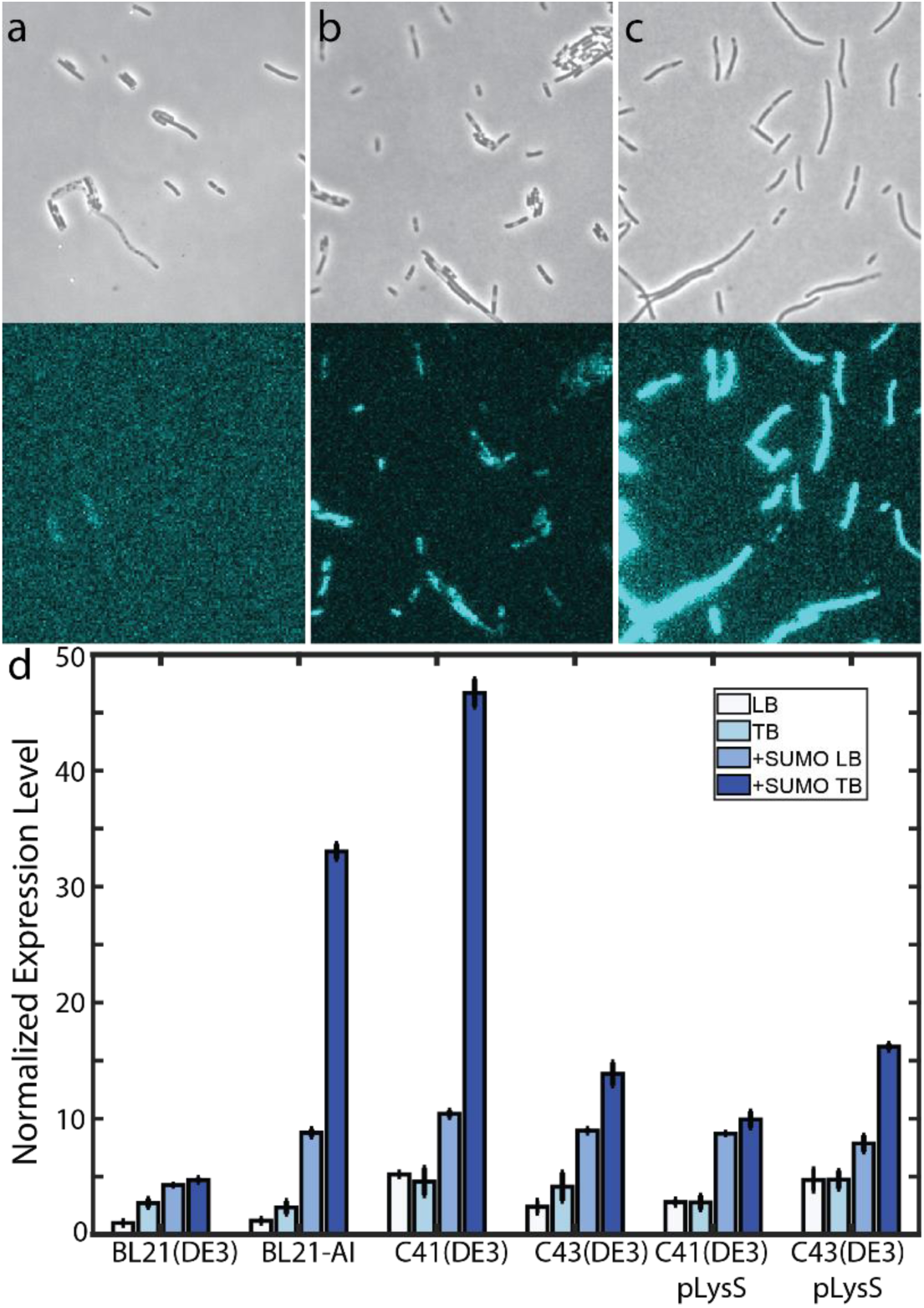
Membrane protein expression in *E. coli.* **(a-c)** Fluorescence microscopy of M-mCerulean3 expression in TB medium, with phase contrast image shown above and mCerulean3 fluorescence channel shown below. (**a**) M-mCerulean3 expressed in BL21(DE3). (**b**) SUMO-M-mCerulean3 expressed in BL21(DE3). (**c**) SUMO-M-mCerulean3 expressed in C41(DE3). All strains were imaged with identical microscope and camera settings, and images are displayed with identical lookup tables (LUTs). (**d**) Quantitative comparison of M-mCerulean3 expression in various *E. coli* strains and media (white – LB; light blue – TB; SUMO-M-mCerulean3 in LB – medium blue; SUMO-M-mCerulean3 in TB – dark blue. Expression levels are normalized to expression of M-mCerulean3 in BL21(DE3).

Previous reports have demonstrated that expression of M protein from the original SARS-CoV virus could be significantly enhanced by N-terminal fusion of M with a Small Ubiquitin-related Modifier (SUMO) tag ^7^. Following this same strategy resulted in enhancement of expression by ∼4-5x (Fig. 1b,d). We next attempted to further optimize expression by quantifying the level of expression of M-mCerulean3 and SUMO-M-mCerulean3 in various strains of *E. coli*, including BL21-AI and the Walker strains, C41(DE3) and C43(DE3), which have been specifically selected for high expression of membrane-bound and other difficult to express proteins. We measured expression in these strains grown in both lysogeny broth (LB) and Terrific Broth (TB), finding that expression of SUMO-M-mCerulean3 in C41(DE3) grown in TB to be ∼45 – 50x higher than that of the standard use of BL21(DE3) grown in LB.

Purification of SUMO-M from C41(DE3) grown in TB using standard immobilized metal affinity chromatography (IMAC) resulted in extremely high yields, typically on the order of 50 – 100 mg per liter of culture. Following purification, we refolded the protein by dialysis and removed the 6xHis tag and SUMO tag by digestion with Ulp1 SUMO protease, resulting in pure, full length, native M protein. We quantified the purity by SDS-PAGE, western blot, and matrix-assisted laser desorption/ionization (MALDI) (Fig. 2) and quantified the endotoxin levels by LAL endotoxin assay, finding an endotoxin content of 0.35 ± 0.05 EU/mg. SDS-PAGE (Fig. 2a) and MALDI mass spectrometry (Fig. 2d) show residual cleaved 6xHis-SUMO tag (12.4 kDa), which was subsequently removed by further IMAC purification with nickel resin. Furthermore, western blot shows (Fig. 2b,c), in addition to a large band at the predicted molecular weight of 25.2 kDa for M monomers, a large streak of high molecular weight multimers and aggregates as well as a small amount of lower molecular weight products that may be the result of degradation. However, as revealed by SDS-PAGE (Fig. 2a), western blot (Fig. 2b,c), and MALDI mass spectrometry (Fig. 2d), the majority of detectable products are M monomer and aggregates thereof.

**Figure 2.**
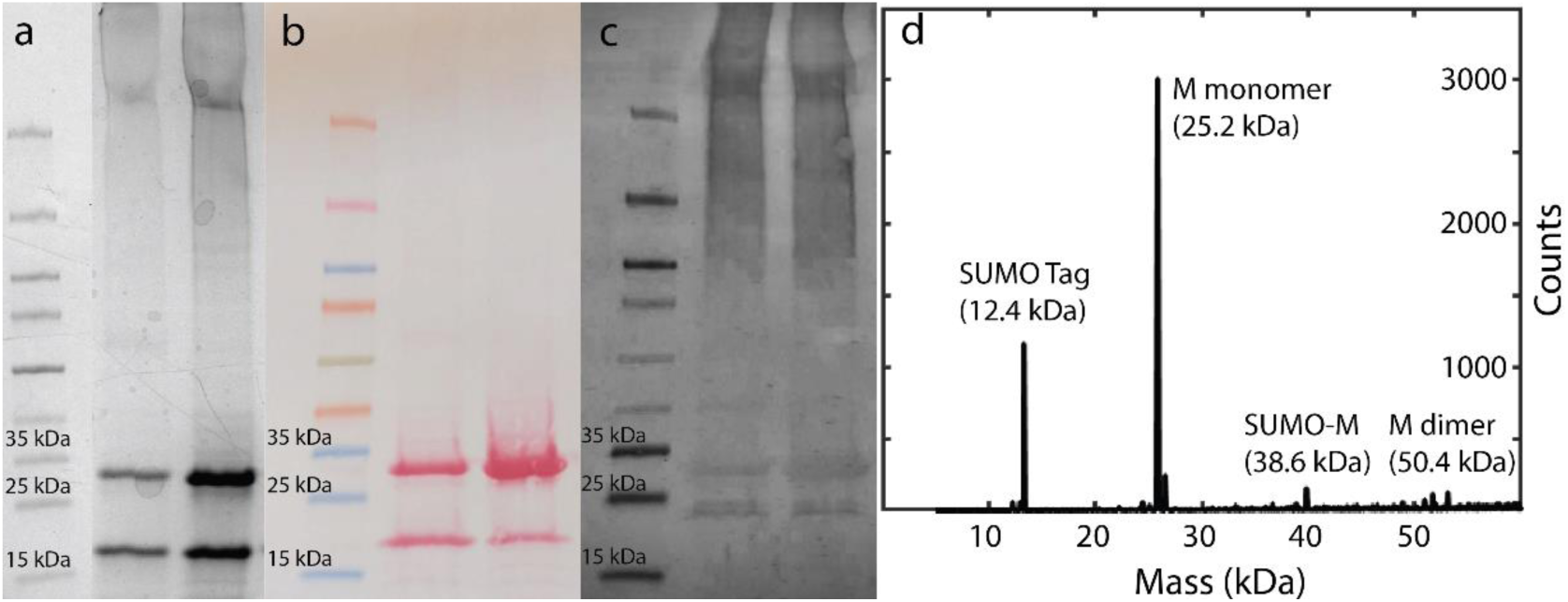
Verification of purified M protein. (**a**) SDS-PAGE of purified M sample. Lane 1: ladder; Lane 2: After digestion with 0.5 mg SUMO protease per mg M; Lane 3: digestion with 2x amount SUMO protease. (**b**) Poinceau all-protein stain of protein transferred to blot from PAGE gel. Lanes same as (a). (**c**) Western blot of (a). (**d**) MALDI mass spectrum of purified M sample.

### Cryo-EM of lipid bilayer membrane reconstituted with M protein in large unilamellar vesicles (LUVs)

The procedure for the M protein insertion is described in Methods and in the Supplemental information (Fig. S1). We used Cryo-EM and the AFM to verify the presence of the inserted M protein and its orientation below. Cryo-EM studies of LUVs with and without M protein were done to confirm the presence of M proteins in the bilayer membranes. A representative Cryo-EM image of the M protein embedded in bilayer vesicles is shown in Fig. 3a, with the schematic of LUVs with (Fig. 3b) and without (Fig. 3c) M protein embedded. Figures 3d-f show the typical 2D class average images of Cryo-EM observations from LUVs reconstituted with M protein at a protein-to-lipid mass ratio of 0.015. Two types of 2D class averages were observed in the Cryo-EM. Figures 3d,e show the M protein as bright spots on the bilayer corresponding to higher electron density of the M protein embedded in the membrane, while in the image in Fig. 3f there is no M protein embedded. Most frequently observed spots were ∼ 4-5 nm wide, which can be associated with single dimers or possibly some higher order oligomers. We also observed that these spots corresponding to the M protein show a higher intensity on the inner leaflet of the vesicle, indicating they are likely inserted with the larger and denser C-terminal facing the inside of vesicles. In contrast with regions reconstituted with M proteins, the regions without M proteins show homogeneous intensity from the corresponding uniform electron density throughout the bilayer membrane. In the bottom row as a negative control, blank vesicle samples without M protein were also imaged with Cryo-EM. Fig. 3g-i show typical 2D class average images with a homogenous electron density observed for the LUVs without M protein. Lastly, Fig. 3j shows a simulation of these Cryo-EM micrographs with M protein calculated from the molecular dynamics simulations of the short form of M protein embedded in a membrane by projecting the atom number density onto a plane perpendicular to the surface of the membrane. Two different viewing angles are averaged over in creating this simulated image, and lower resolution density maps for both the long and short forms shown in Fig. S5.

**Figure 3.**
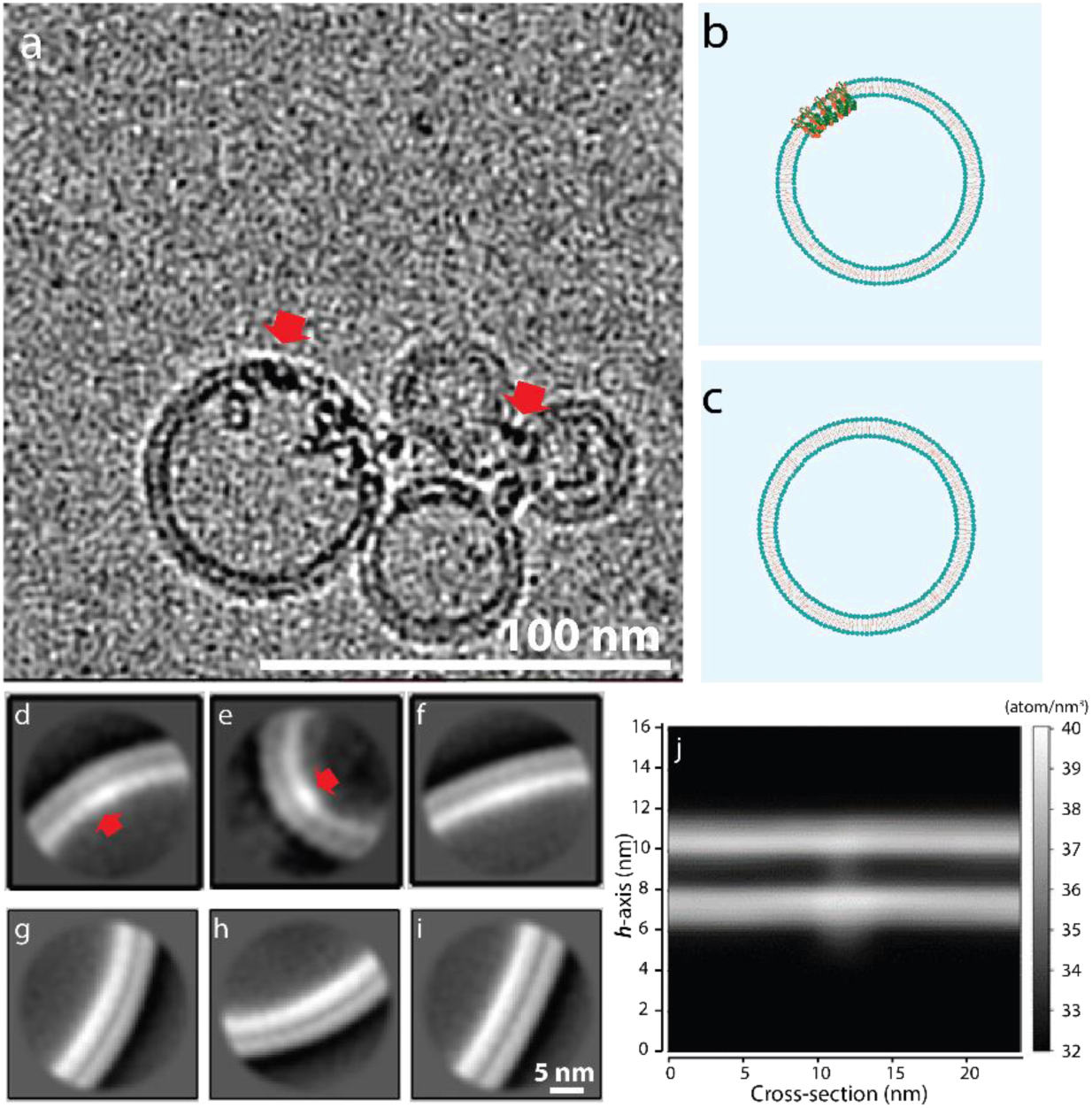
Typical CryoEM micrographs of LUVs reconstituted with M proteins. (**a**) Red arrows indicate M proteins inserted in the vesicle membrane. (**b**) and (**c**) - schematics of LUVs with and without M reconstitution, respectively. (**d**) Representative 2D class average image of areas without M protein in reconstituted LUVs. (**e**) and (**f**) – two most representative CryoEM 2D class average images from LUV bilayer membranes reconstituted with M protein mass ratio, R_m,M_/L = 0.015. Arrows: location of M protein as bright spots. (**g**-**i**) 2D class average CryoEM images of blank LUV bilayer membrane. Scale bar = 5.0 nm. (**j**) Atom number density of the short conformation M protein embedded in a bilayer membrane obtained from all atom MD.

### AFM topography of SBL reconstituted with embedded M protein

As presented above the Cryo-EM confirmed the presence of the M protein in the lipid bilayer. The small molecular weight of 50.2kD for the dimer limits high resolution analysis of the M protein-lipid membrane interaction. The AFM with the height resolution of 0.01nm and a lateral resolution 0.8 nm is ideally suited for understanding the inserted protein morphology, orientation, oligomerization at larger concentration and the protein-lipid bilayer interaction. Figure 4a shows a 1 µm x 1 µm AFM topography of a typical endoplasmic reticulum-Golgi intermediate compartment (ERGIC) bilayer membrane generated from LUVs without M protein reconstitution. The height profile along the green dashed line is shown in the panel below. The SBL on the mica substrate is smooth with an RMS roughness measured to be 0.3 nm. Figure 4b shows the typical topography of a 1 µm x 1 µm surface of a SBL generated from LUVs reconstituted with M protein at a protein-to-lipid mass ratio of 0.01. In contrast to the smooth surface of the SBL in Fig. 4a, scattered single particle protrusions were observed in the AFM image. These can be associated with the embedded protein in the SBL. The line-profile (green dashed line) passing through one of the M proteins shows that the height of the feature above the membrane is 2.4 nm, consistent with the C-terminal height of M protein from literature ^3, 4^ and simulations presented below. Figures 4c,d show the SBL from LUVs reconstituted at higher protein-to-lipid mass ratios of 0.015 and 0.02 respectively. Figure 4c topography shows a combination of scattered single particles and patches, while patches are mostly observed in Fig. 4d. The line profiles show that the patches have the same height as the single particles associated with the M protein. Figure 4e shows the percent of protein-occupied area in the SBL membrane as a function of protein-to-lipid ratio. The area occupied by the protein is calculated from the sum of the single protein and patch areas. The M protein-occupied area increases from 0.3% to 6.2% and 17.1%, as the protein ratio increased from 0.01 to 0.015 and 0.02, respectively.

**Figure 4.**
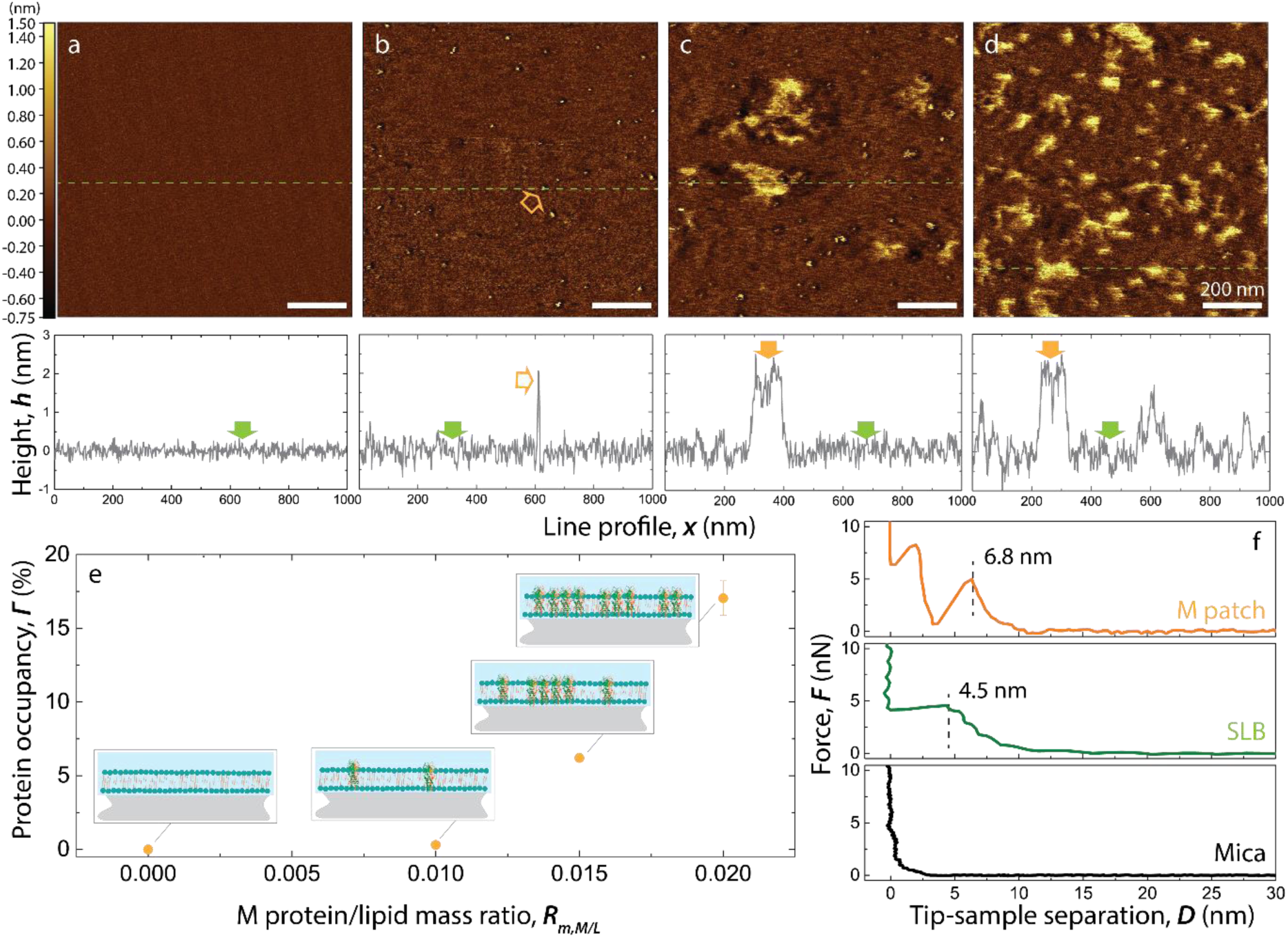
AFM height image of supported bilayer (SBL) reconstituted with M protein at increasing protein to lipid mass ratios, R_m,M/L_, imaged on mica substrates. (**a**-**d**) Height images and representative single line profiles (green dotted lines in images) collected from SBL fabricated from large unilamellar vesicles that were reconstituted with M protein at R_m,M/L_ = 0, 0.01, 0.015 and 0.02. (**e**) Area occupation percentage of M protein in SBL as a function of R_m,M/L_ calculated from A-D. (**f**) Representative force profiles collected from pristine mica surface (black), SBL (green) and M protein (orange) patches in lipid bilayer, the force profiles were collected during tip engagement.

In addition to examining the planar surface features of the SBL with and without M protein we also characterized the depth profile using AFM indentation studies. Figure 4f shows a typical indentation force profile obtained from three different samples: bare mica in the bottom panel; SBL without and with M protein (concentration ratio of *R* = 0.02) in the middle and top panels respectively. In all three panels the horizontal axis is the separation distance between the AFM probe tip and the bare mica surface. The bottom panel shows the typical indentation force curve of the bare mica surface in solution with the tip-rigid surface contact at zero followed by a sharp increase in force by the cantilever bending. The middle panel corresponds to the SBL membrane without M protein. Here, the AFM tip comes in contact with the hydration layer at a separation around 7.5 nm with the applied force leading to deformation of the membrane. With increasing applied force, the tip punctures the membrane surface at *D* = 4.5 ±0.1 nm. This corresponds to the membrane thickness without M protein. With further increase in force the AFM tip penetrates the membrane until it contacts the rigid mica at 0 nm. The top panel shows the indentation force profile obtained from the patch-like structures from the M protein on the SBL. The force showed a discontinuity at 6.1 nm which corresponds to the tip-protein surface interaction. Large forces lead to penetration of the tip into the M protein patch. The discontinuity at 3.2 nm indicates potential disruption of a specific strong interaction region between neighboring M proteins. From the slope of the approach curves in Fig. 4f, the Young’s Modulus can be calculated to be 9.5 MPa on the lipid bilayer and 41.5 MPa on the protein patches respectively. Figure S2 shows the average elastic moduli extracted from nanoindentation force profiles for SLB and the M protein patches. The difference in the Young’s Modulus indicates that the patches are indeed a different type of material from the lipid bilayer.

### Formation of M protein aggregate patches in the Supported Bilayer (SBL)

From Fig. 4b we observed that M proteins in SBL start as single M protein assemblies at a M protein to lipid mass concentration *R* = 0.01. From the size (length, width, and height) they can be identified as M protein dimers. With the increase of M protein concentration in the SBL the dimers aggregate to form patches of M proteins as observed in Figs. 4c and d. As seen in Fig. 4e the area of the patches grows nonlinearly with the M protein concentration. The M protein-occupied area increases from 0.3% to 6.2% and 17.1%, as the protein to lipid mass ratio R_m,M/L_ increased from 0.01 to 0.015 and 0.02. Thus, the M protein incorporation into the membrane is facilitated by the presence of M proteins already in the membrane. Notice that the height of patch-like assemblies is the same as the isolated single M protein dimers observed at the lower M concentration. This indicates that the M proteins assemble in a side-by-side manner inside the bilayer, as demonstrated by schematics shown in Fig. 4e; these patches are higher order oligomers.

### Morphological statistics of the isolated M protein particle

We next characterized the properties of single M protein particles embedded in the SBL for protein to lipid mass ratio, R_m,M/L_=0.01 (Fig. 5). In Fig. 5a we show a high resolution 250 x 250 nm surface morphology of a typical single isolated M protein at the concentration ratio of R_m,M/L_= 0.01. In Fig. 5b the characterization parameters the Length (L) and Width (W) are shown. The boundary of the protein particle is identified as an edge higher than the planar membrane height. The planar membrane height is found by averaging the membrane height far away from the particle (gray region, Fig. 5b). The particle thus identified is shown in red in Fig. 5b. The Length (L) of the particle is defined as the maximum distance between any two points on the perimeter of the protein particle. The particle Width (W) is defined as the maximum distance between two points on the perimeter, in the direction perpendicular to length. The particle Height (H) is the height of the highest point on the protein surface. For comparison, Fig. 5c shows the final frame of an all-atom molecular dynamics simulation for the short conformation M protein embedded in a multicomponent membrane. The analysis of the AFM data is repeated for 214 samples and the histogram of the results for L, W, H and the ratio L/W are shown in Fig. 5d-g. Corresponding size classifications from molecular dynamics simulations are shown in Fig. 5d-g, averaged over the last 100 ns of the simulation, through the dotted purple lines (see Fig. S9-S11 for more information). Protein length and width were obtained by fitting an ellipse to the protein, while height was based off the difference between the top of the M protein C-terminal and the phospholipid heads adjacent to the protein. The histogram in Fig. 5d shows a bimodal distribution which when fit to two normal distributions lead to mean particle lengths, L_1_ = 4.8 ± 2.0 nm and L_2_ = 9.4 ± 3.1 nm. Figure 5e shows the width distribution, again seen to be bimodal, which when fit to two normal distributions leads to mean values of W_1_ = 2.3 ± 1.7 nm and another at W_2_ = 5.7 ± 1.4 nm. In Fig. 5f the aspect ratio ‘L/W’ again shows two normal distributions, one at L/W_1_ = 1.9 ± 0.5, the other at L/W_2_ = 3.1 ± 0.6. Figure 5g shows the histogram obtained from the height of the M protein particles. The values can be fit to one normal distribution with a mean height h = 2.4 ± 0.7 nm.

**Figure 5.**
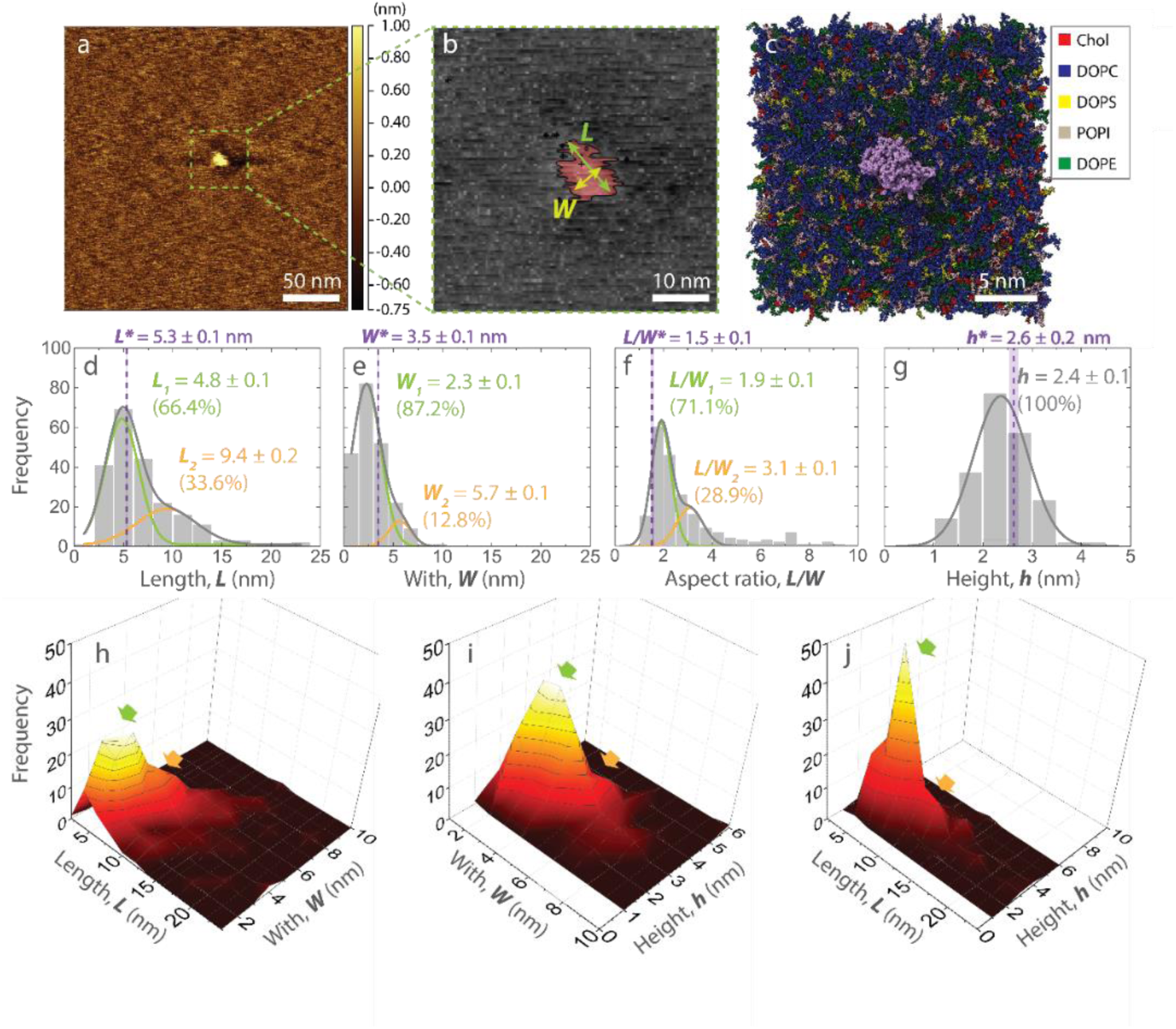
Size and morphology statistics of M protein quantified from AFM images of SBL. (**a**) Representative AFM height image of individual M protein reconstituted in SBL at R_m,M/L_ = 0.01. (**b**) Zoomed-in image of (a), highlighting the protein with red color and lipid bilayer with gray color. The protruded portion of a single M protein particle above lipid bilayer (left) is detected by setting the height of top surface of SBL as a threshold and taking the above portion. (**c**) Top view of the short conformation M protein (C-terminal in purple) embedded in a multicomponent membrane 1 µs into an all atom MD simulation. Dimensional parameters: length, L, width, W, and height, h, were measured for detected protein particles. (**d**-**g**) Histograms of L, W, L/W, and h of protein protruding above lipid membrane. Solid lines are Gaussian fits. Dotted vertical lines mark corresponding values from MD simulation for short-form M protein reconstituted with C-terminal exposed above SBL. (**h**-**j**) 3D histograms each showing combined statistical survey of two out of the three dimensions. Green and orange arrows indicate dimer and higher order of oligomers.

### M proteins are confirmed to be inserted in the bilayer membrane

High resolution Cryo-EM studies have shown that the SARS-CoV-2 membrane protein dimers exist in a conformational equilibrium between the long form and the short form, which are 7.2 nm and 8.6 nm in height, respectively ^3^. Comparing these heights with Fig. 5f, AFM shows the height of exposed protein particles is much shorter (2.4 nm). This is in agreement with the all atom MD simulations, where the height ranges from 2.6 nm to 2.8 nm depending on the protein conformation (short/long). The slight differences between AFM and MD can be attributed to the protein deforming slightly as the AFM tip makes contact. Meanwhile, the lateral dimensions of isolated protein particles observed from AFM in Fig. 5c,d are very similar to the literature reported values of 5.0 nm and 5.7 nm for long and short form respectively ^3^. A much smaller height and similar lateral dimensions indicate that the M proteins are inserted into the bilayer membranes instead of absorbing on top of the membranes. This is also supported by our Cryo-EM observations in Fig. 3b,c. The membrane reconstituted with M protein shows a higher electron density compared with blank membrane. This indicates that a large portion of the protein is buried inside the bilayer membrane.

### Orientation of reconstituted M protein has C terminal facing the vesicle interior

In Fig. 3b,c, we observed that the bright spots corresponding to reconstituted proteins show a higher electron density in the inner leaflets. This indicates that the C terminus, which is larger than the N terminus and contains a higher density of electron-rich “heavy” atoms, is inserted facing the inside of the vesicles. This is further confirmed in AFM measurements and also matches the simulated Cryo-EM image from MD in Fig. 3j. This stems from averaging the protein over different angles, since the N-terminus of the protein on the outer leaflet ranges from small to large arc length along the membrane. Alternatively, the C-terminal domain of the protein along the inner leaflet is largely independent of the viewing angle and will have greater arc length than the N-terminal domain. In Fig. 5f, the height of reconstituted M protein exposed above bilayer membrane show a single distribution at 2.4 nm, which is consistent with C terminal size observed in high resolution Cryo-EM images and our simulation predictions, indicating that all M proteins in the SBLs observed in AFM are oriented with their C terminals facing the AFM probe. From these results we can deduce that the M proteins were reconstituted in LUVs with their C-terminal encapsulated inside the vesicles. This is because, during the formation of the SBL on mica, the LUVs adsorb to the mica surface with their outer leaflets. Subsequently the LUVs rupture and expose their inner leaflets to the AFM probe.

### Reconstituted M protein shows dimensions and membrane thickness consistent with the short form

In Figs. 5h-j the three-dimensional histograms analyze the relationships between protein length (*L*) and width (*W*), width and height, as well as length and height (*h*), respectively. In Fig. 5h from the *L* and *W*, we can observe two distinct populations with their individual lengths and widths. From Fig. 5i plot of the *L* and *H* we can observe that both *L* populations have the same height above the membrane. From Fig. 5j the same is observed for the two width populations. Combining the above information with results shown in the previous two-dimensional histograms (Figs. 5c-f), the most frequently observed M protein population in the SBL have a length, width and height of 4.8 ± 0.1 nm, 2.3 ± 0.1 nm and 2.4 ± 0.1 nm, respectively. These dimensions show good agreement with the short form of M protein dimer both from published Cryo-EM study of M protein in detergent micelles^11^ and our simulation predictions (shown with the purple lines in Figs. 5d-g). The slight discrepancy between the short conformation MD dimensions and AFM results likely stem from the original MD protein structures not including every residue and deformations in the protein from the AFM tip. Furthermore, the membrane thinning around the protein described in Fig. 5b agrees with the observations from Neuman et al. for SARS-CoV, where the short conformation is found in thinner regions of membrane as opposed to the long form. Additionally, the AFM determined thinning profile matches the MD short form thinning profile, as opposed to the long form (long form shown in Fig. S4). The second population observed from above has a much larger size with a length, width and height of 9.4 ± 0.2 nm, 5.7 ± 0.1 nm, and 2.4 ± 0.1 nm, respectively. These dimensions correspond to that of a multimer of M protein dimers. Fig. 5g shows the schematics of top view of individual dimer in the short form, as well as possible oligomer configurations corresponding to the observed second population.

### Reduction of membrane thickness around M protein

Figure 6a shows a typical isolated M protein particle in a magnified view, with color scale corresponding to the membrane thickness shown to the right. We observed that the membrane thickness in proximity to the single M protein particle is reduced. The height of the membrane around each protein particle is measured along two perpendicular axes, shown in Fig. 6a as dashed green lines. A similar view from above of the M protein short conformation obtained through all atom molecular dynamics can be seen in Fig. 6b, where red regions represent a thicker membrane and blue thinner. The circular cross section of the protein shown in gold stems from its rotation shown in Fig. S8. Figure 6c shows an exaggerated diagram of the membrane thinning near the protein and a corresponding side view of the thickness after 1 µs of molecular dynamics simulation is displayed in Fig. 6d. The change in membrane height from the edge of the protein (0 nm) along the dashed lines for 25 such proteins is shown in Fig. 6e. Additionally, the yellow points represent shifted membrane thickness values from Fig. 6d, based on the edge of the protein. The mean change in height with distance from the protein edge is shown by the solid green line. We observed that the bilayer membrane is compressed until the height appears to saturate 12 nm from the edge of the protein. The maximum membrane compression at the edge of the protein has a mean value of 0.5 nm.

**Figure 6.**
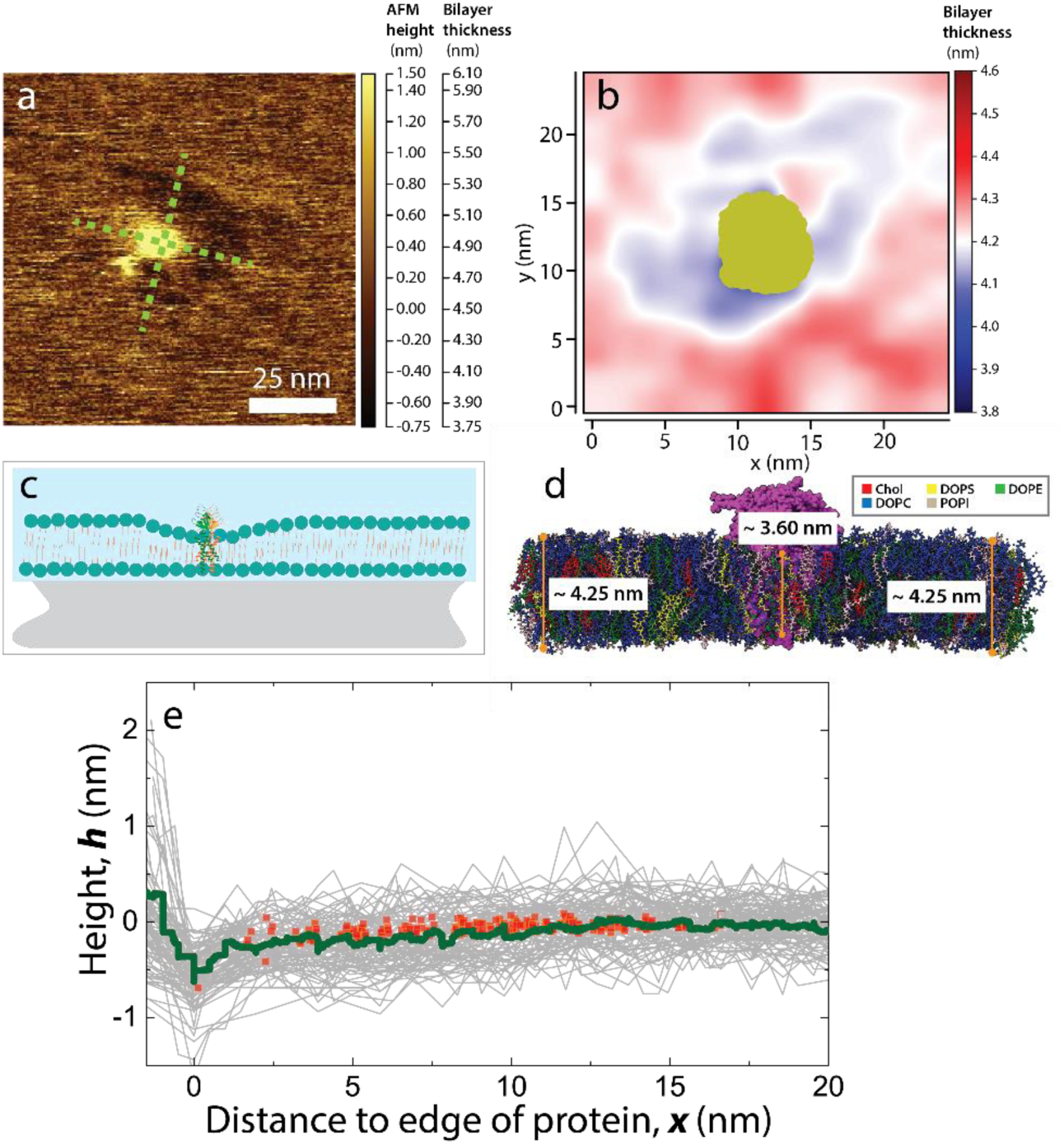
Analysis of membrane thickness around individual M protein particles at R_m,M/L_ = 0.01 and through MD simulation. (**a**) Example of individual M protein particles reconstituted in SLB. Height profiles were taken along the green dotted lines starting from the center of the protein. (**b**) View from above of membrane thickness in nm from 500 ns – 1 µs of an all-atom MD simulation of the short conformation M protein embedded in a multicomponent membrane. (**c**) Schematics of membrane thinning around M protein particles. (**d**) Side view of the MD simulation along the y-axis, with the protein in purple, where average thickness is shown dependent on distance from the protein. (**e**) Solid green line shows the experimentally measured averaged height profile of membrane around individual M protein particles. Inset shows around 100 height profiles obtained from 25 individual protein particles. Here *x* = 0 nm is defined by the boundary between the M protein particle and lipid membrane. The yellow points represent vertically shifted values of membrane thickness obtained through MD shown in (d).

### Membrane compression driven by hydrophobic mismatch of protein and lipid membrane, M protein aggregation and spontaneous membrane curvature

The membrane thickness reduction around M protein observed in Fig. 6 is likely due to the height mismatch between the hydrophobic transmembrane domain and the thickness of the lipid bilayer. From the literature ^8^, and confirmed with our all-atom MD (Fig. S9d), the transmembrane region of the M protein dimer is around 4 nm. The experimentally measured thickness of the lipid bilayer is 4.5 nm as shown in Fig. 4f. Thus, the protein hydrophobic region is ∼0.5 nm smaller than the membrane thickness. This hydrophobic mismatch leads to compression of the bilayer in the immediate proximity of the M protein. In Fig. 6b the change in the thickness of the membrane is shown as a function of distance from the edge of the protein. The maximum compression at the edge of the protein is 0.5 nm seen through AFM. Measurable compression extends to a distance of 12 nm from the edge of the protein. From all atom MD simulations of the M protein dimer embedded in a membrane, this thinning profile agrees with that seen for the short conformation of the M protein. Thinning/thickening results for the long conformation can be seen in Fig. S6, although the protein deviates from its initial structure quickly. A more thorough discussion of these results can be seen in the Supplementary Information. The thinning of the membrane in the vicinity of the protein results in an energetic cost due to bend and tilt deformations in the lipid layers, which gives rise to an effective line tension ^9^. Using our measurements of the mechanical moduli of the membrane, informed by estimates from literature ^10^ and the magnitude of the thinning, we can estimate the effective line tension to be about 0.5 pN (see Supplementary Information for details). This value is consistent with estimates and measurements of line tension for lipid domains with comparable height mismatches ^9, 11^, which control the growth and size of lipid ordered domains ^11, 12^, suggesting that the line tension can facilitate the aggregation of M into patches. Our coarse-grained simulation shows that this patch formation leads to the spontaneous curvature of the membrane. However, the attractive interaction between endodomains is essential for the membrane budding (Fig. 6). Interestingly, our ERGIC simulation reveals that N proteins and RNA can effectively introduce membrane curvature as well, providing insight into the mechanism underlying virus budding and encapsulation (Fig. S12)

### Coarse-grained Molecular Dynamics Simulation of M protein-induced budding

To illustrate how the inserted M proteins can give rise to virus budding, we further perform coarse-grained simulations by modeling M proteins and a flat lipid bilayer as shown in Figure 7a. Each M protein is built from three hard particles (M1, M2, and M3), where M1 and M2 correspond to the transmembrane domain, and M3 denotes the endodomain. The membrane is modeled as a triangular lattice with hard particles (M) occupying the vertices, on account of the lipids’ incompressibility. Since most of experiments and all-atom simulations in this paper are performed on the flat membrane, we also employ a flat membrane to investigate the role of membrane protein on curvature. The simulation results are shown in Figures 7b,c, where M proteins interact with each other through Lennard-Jones potentials. Figure 7b shows that transmembrane domain interaction (M2-M2) cannot give rise to M protein-induced membrane curvature leading to the membrane budding even if the budding particle has lower energy. This is mainly due to the presence of the energy barrier required to bend the membrane. We find that an attractive endodomain interaction is necessary under all the conditions examined to overcome the bending energy penalties. Figure 7c shows that as we increase the M protein endodomain interaction (M3-M3), the membrane with the embedded M proteins can bud and form a spherical shell. We note that N proteins and RNA can also effectively provide the M protein endodomain interactions and help M proteins bud ^13^, see Fig. S12. The budding prerequisites could indicate the mechanism of how coronavirus can encapsidate N proteins and RNA complex into an infective particle ^13^.

**Figure 7.**
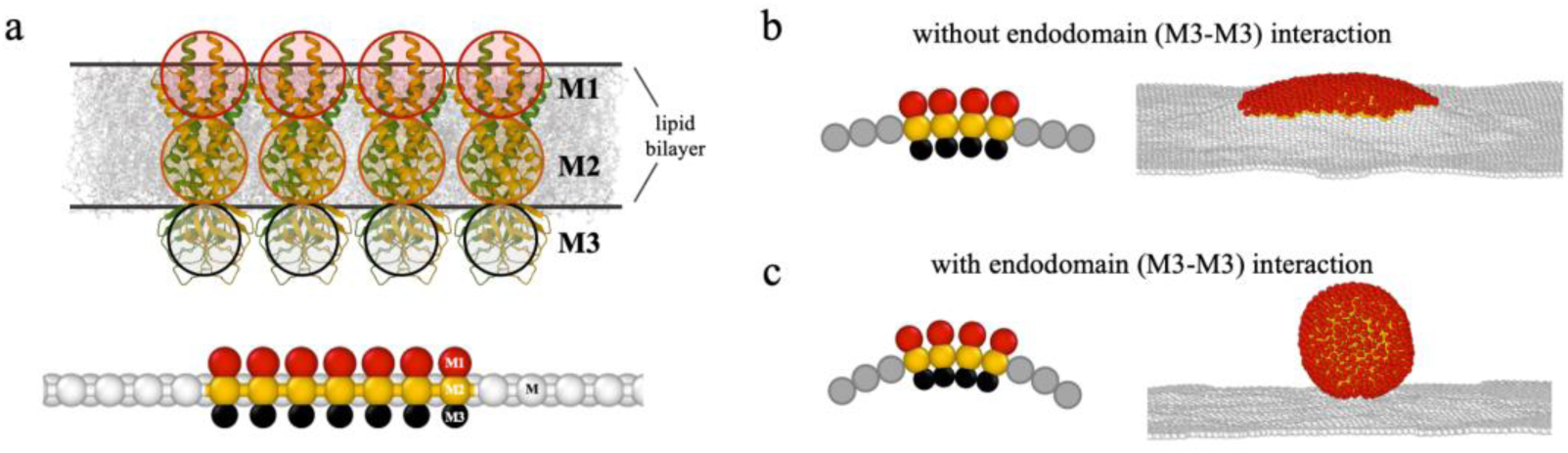
Coarse-grained simulations of M proteins embedded in a flat membrane. (**a**) Schematic illustration of M proteins embedded in lipid bilayers. The upper figure shows the actual size of the lipid bilayer and M proteins (PDB 7VGR ^3^), and the lower figure shows the coarse-grained model used in the simulation. (**b**) Schematic figure (left) and simulation snapshot (right) for the case that the M proteins have no endodomain interactions (*ϵ*_*M*3*M*3_ = 0). The M proteins aggregate and form a small bump due to the line tension, but no further budding is observed in the simulation. (**c**) Schematic figure (left) and simulation snapshot (right) for the case that M proteins have endodomain interactions (*ϵ*_*M*3*M*3_ = 8*k*_*B*_*T*). The interaction helps M proteins overcome the energy barrier and the membrane buds into a spherical shell. For each M protein, the domain particles have a diameter size of 1.0*a* (M1), 1.0*a* (M2), and 0.8*a* (M3), where *a* is the system length unit corresponding to 3*nm*. For both simulations, the transmembrane domain M2 interacts with M2 with strength *ϵ*_*M*2*M*2_ = 4*k*_*B*_*T*. The membrane is represented as a triangular lattice where rigid particles (M) occupy the vertices due to the incompressibility of lipids membrane. The M particles interacts with M with strength *ϵ*_*MM*_ = 4*k*_*B*_*T*. The rest of particles have excluded volume repulsion defined through a cutoff Lennard-Jones potential.

## DISCUSSION

While Membrane (M) protein is the most abundant protein present in the SARS-CoV-2 particles, its detailed characterization has proven challenging due to difficulties with expression and purification. We present here a method for the purification of large amounts of SARS-CoV-2 Membrane (M) protein from *E. coli*. The advantage of *E. coli* as an expression system is the simplicity and low cost of production and purification resulting in extremely high yields. However, it should be kept in mind that *E. coli* is unable to perform post-translational modifications, such as glycoslylation. Like the SARS-CoV-2 S and E proteins, M protein is predicted to be post-transcriptionally glycosylated ^14^, and eight possible sites of N-linked glycosylations have been predicted using bioinformatics comparisons with M proteins in other viruses in the Covid family ^15^, although it is not yet clear the extent or function of this modification. However, it is known that for the SARS-CoV virus these modifications are not required for viral assembly or protein-protein interactions ^16–19^.

Using the purified M protein in combination with a phospholipid composition mimicking that of the ERGIC cellular compartment where viruses are assembled *in vivo*, we used CryoEM and Atomic Force Microscopy (AFM) to study M protein embedded within a lipid membrane and its interactions with the membrane. We observe that M protein is inserted into the membrane with its C-terminus on the interior of vesicles, consistent with budding *in vivo*, and that interactions of M with the membrane result in a local reduction in membrane thickness. We then use all-atom molecular dynamics simulations to show membrane thinning consistent with experiment and the propensity for the M proteins to induce membrane curvature. We estimate the line tension resulting from membrane thickness reduction and suggest that this could provide a fundamental mechanism driving M protein aggregation. Taken together with potential membrane curvature induction, our coarse-grained simulations show that these features can result in viral assembly and budding.

## METHODS

### Resource Availability

#### Lead Contact

Further information and requests for resources and reagents should be directed to and will be fulfilled by the lead contact, Thomas E. Kuhlman. All data reported in this paper will be shared by the lead contact upon request.

#### Data and Code Availability

Any additional information required to reanalyze the data reported in this paper is available from the lead contact upon request.

#### Materials Availability

Plasmids and strains used in this study are available from the lead contact upon request.

### Strains and Media

For protein expression and purification experiments we employed the commonly used protein expression strains BL21(DE3) (ThermoFisher Scientific) ^20, 21^, BL21-AI (ThermoFisher Scientific), C41(DE3) ^22^, C43(DE3) ^22^, C41(DE3) pLysS ^22, 23^, and C43(DE3) pLysS ^22, 23^. Cells were grown in Lysogeny Broth - Miller (LB-Miller) [10g tryptone (Bacto, ThermoFisher Scientific), 5g yeast extract (Bacto, ThermoFisher Scientific), 10g NaCl (Millipore Sigma), and ultrapure H_2_O (Milli-Q IQ 7000, Millipore Sigma) to one liter medium ^24, 25^] or Modified Terrific Broth (MTB) [20g tryptone, 24g yeast extract, 4mL glycerol (Fisher Scientific), 0.017 M KH_2_PO_4_ (Millipore Sigma), 0.072 M K_2_HPO_4_ (Millipore Sigma), and ultrapure H_2_O to one liter medium ^26^].

### Plasmid Construction

Plasmids encoding and expressing M with an N-terminal 6xHis tag and with and without SUMO tag fusions were designed in VectorNTI software (ThermoFisher Scientific) and synthesized *de novo* by GENEWIZ/Azenta Gene Synthesis service. Constructs were synthesized to be expressed from a T7*lac* promoter ^27^ with a consensus Shine Dalgarno ribosomal binding site and were cloned into plasmid pUC57-kan-GW. A fluorescently tagged variant, M-mCerulean3 ^28^ was designed and produced similarly with C-terminal fluorescent fusion.

### Cell Growth and Protein Expression and Purification

Cells were grown identically for all experiments. First, a seed culture was started by inoculation of 25 mL medium containing 25 µg/ml kanamycin in a baffled 125 ml Erlenmeyer flask from freezer stock and allowed to grow overnight at 37 °C in a C76 shaking water bath (New Brunswick Scientific). This culture was used to inoculate 1 L of medium containing 25 µg/ml kanamycin in a 2 L baffled Erlenmeyer flask and grown at 37 °C in a C76 shaking water bath until the optical density at 600 nm (OD_600_, SmartSpec Plus spectrophotometer, Bio-Rad) reached ∼0.5 – 0.6. At this point, the flask was removed from the water bath and transferred to a shaker (Dura-Shaker, VWR) at room temperature. After cooling to room temp, the appropriate concentration of inducer to effect full induction was added [0.4% w/v L-arabinose for BL21-AI or 1 mM isopropyl β-D-1-thiogalactopyranoside (IPTG) for all others], and the culture allowed to grow overnight to saturation.

### Quantification of Protein Expression and Microscopy

To quantify expression of M protein in various *E. coli* strains and media, cells transformed with the indicated plasmids expressing variants of M-mCerulean3 were grown as described above. 2 mL of culture was then collected and centrifuged (10,000g Eppendorf 5424) and the supernatant medium aspirated away. Cells were then washed with phosphate buffered saline (PBS) three times, and finally resuspended in fresh PBS. Cells were diluted with PBS such that OD_600_ < 1.0 and distributed to the wells of a 48 well plate (Corning Costar). The plate was placed in a CLARIOstar microplate reader (BMG Labtech) and the OD_600_ and mCerulean3 fluorescence measured (Excitation at 430/20 nm, emission at 480/20 nm). Expression levels reported in Figure 1 are calculated as mCerulean3 fluorescence divided by OD_600_. Each measurement was performed with three technical replicates over three experimental/biological replicates.

Fluorescence microscopy was performed on a Nikon Ti-E inverted fluorescence microscope with an Andor DU-897 Ultra camera. For imaging of M-mCerulean3 fluorescence, cells were imaged with a 100x TIRF objective and illuminated with highly inclined 457 nm laser light from a Milles Griot argon laser fitted on a Nikon LU-4A laser launch and using a Chroma filter cube with Z457/10x excitation and ET485/30m emission filters.

### Protein Purification

After growth to saturation as described above, the 1 L culture was spun down for 30 min at 4,300g in an Eppendorf 5910Ri refrigerated centrifuge maintained at 4 °C and the resulting supernatant discarded. Cells were resuspended in 10 mL lysis buffer (50 mM NaH2PO4, 300 mM NaCl, 10 mM imidazole, pH = 8.0) with 1 mg/ml lysozyme (Millipore Sigma) and 50 U/ml benzonase nuclease (Millipore Sigma) and incubated at 4 °C for 30 minutes on a nutating shaker. The cells were flash frozen in liquid nitrogen, followed by incubation in a room temperature water bath until thawed. This slurry was then centrifuged at 10,000 g at 4 °C for 30 minutes. The supernatant clarified cell extract (soluble fraction) was collected and stored at −80 °C until purification. The pelleted cell debris was then washed three times in inclusion body wash buffer [PBS with 25% w/v sucrose, 5 mM EDTA, and 1% Triton X-100] followed by centrifugation at 20,000 g for 30 minutes at 4 °C. After the final wash, inclusion bodies were resuspended in denaturing lysis buffer [8 M Urea (Millipore Sigma), 10 mM Tris (Fisher Scientific), 100 mM NaH_2_PO_4_ (Millipore Sigma), 50 mM 3-(cyclohexylamino)-1-propanesulfonic acid (CAPS, Millipore Sigma), 0.3% N-lauroyl sarcosine (Millipore Sigma), 1 mM dithiothreitol (DTT, Millipore Sigma), pH = 11.0] with nutation at room temperature for 30 min. This insoluble fraction was then stored at −80 °C until purification.

For purification, samples were processed using a Bio-Rad NGC Quest 10 Plus fast protein liquid chromatography (FPLC) apparatus with a 5 mL EconoFit Nuvia IMAC affinity column (Bio-Rad). To purify protein from the soluble fraction, the column was first equilibrated with Wash Buffer A (50 mM NaH_2_PO_4_, 300 mM NaCl, and 20 mM imidazole). The sample was loaded onto the column and washed with five column volumes of Wash Buffer A. The column was then washed with a linearly increasing mixture of Wash Buffer A mixed with Wash Buffer B (50 mM NaH_2_PO_4_, 300 mM NaCl, and 250 mM imidazole) from 3 – 100% Wash Buffer B composition over ten column volumes. Finally, the column was washed with five column volumes of Wash Buffer B. Throughout, the composition of the eluate was monitored by measuring the absorption at 280 nm, and those samples containing the protein of interest were collected and pooled for further processing. To purify protein from the insoluble fraction the procedure was identical using Denaturing Wash Buffer A (50 mM NaH_2_PO_4_, 300 mM NaCl, 50 mM CAPS, 0.3% N-lauroyl sarcosine, and 20 mM imidazole, pH = 11.0) and Denaturing Wash Buffer B (50 mM NaH_2_PO_4_, 300 mM NaCl, 50 mM CAPS, 0.3% N-lauroyl sarcosine, and 300 mM imidazole, pH = 11.0). All purified SUMO-fused proteins eluted as a single peak.

### Protein Refolding and Dialysis

To refold denatured proteins purified from the insoluble fraction, samples were dialyzed in regenerated cellulose dialysis membrane (3.5 kDa MWCO, Spectra/Por) over two days at 4 °C in 10x volume of refolding buffer (20 mM Tris and 10% glycerol, pH = 8.0) with periodic replacement of buffer at least four times over the course of dialysis.

### Cleavage of SUMO tags

To remove the SUMO tag, purified protein (after refolding, if necessary) was mixed with an appropriate volume of 10x SUMO protease cleavage buffer [500 mM Tris, 2% igepal NP-40 (Millipore Sigma), 1.5 M NaCl, and 10 mM DTT, pH = 8.0] along with an excess of Ulp1 6xHis-SUMO protease previously purified from the *E. coli* soluble fraction as described above. Digests were incubated in a 30 °C water bath overnight.

### Removal of SUMO and SUMO protease for final purification

To remove the cleaved 6xHis-SUMO tags and SUMO protease, protein digests were mixed with 1 mL HisPur Ni-NTA resin (ThermoFisher Scientific) and placed on a rotating mixer maintained at 4 °C overnight. After incubation, the samples were applied to an empty Poly-Prep chromatography column (Bio-Rad) and the purified proteins collected as the flowthrough. If higher concentrations were required for a given application, these samples were concentrated using a Pierce PES protein concentrator with appropriate molecular weight cutoff (MWCO, ThermoFisher Scientific).

### Quantification of yields and endotoxin levels

During purification, absorbance at 280 nm was measured by the NGC Quest 10 Plus optical detector and yields calculated using extinction coefficients estimated using ExPASy ProtParam online tool based on protein sequence ^29^. Estimated concentrations were verified using RC DC protein assay (Bio-Rad). Briefly, 5 µl of protein samples and standards (Protein Standards I and II, Bio-Rad) were added to 25 µl reagent A’ and mixed in the wells of a 96 well plate (Corning Costar). To this, 200 µl reagent B was added to each well, mixed, and allowed to incubate at room temperature. After 15 minutes, the absorbance at 750 nm was measured using a CLARIOstar microplate reader, and the concentrations of the samples determined by comparison to linear regression of 2x dilutions of the protein standards.

Endotoxin levels of purified samples were determined using a ToxinSensor chromogenic LAL endotoxin assay kit (GenScript) according to the manufacturer’s instructions. 16.67 µl of samples, endotoxin standards, and water as a negative control were added to the wells of a 96 well plate (Corning Costar). To this, 16.67 µl of LAL was added to each well and mixed. The plate was incubated at 37 °C for 10 minutes. 16.67 µl reconstituted chromogenic substrate was added, mixed, and incubated at 37 °C for six minutes. Then 83.3 µl color stabilizer 1 was added and mixed, followed by 83.3 µl color stabilizer 2, and finally 83.3 µl color stabilizer 3. The absorbance of each sample was measured at 545 nm in a CLARIOstar microplate reader, and the endotoxin levels of the samples were determined by comparison to linear regression using 2x dilutions of the endotoxin standards.

### SDS PAGE and Western Blots

To perform SDS polyacrylamide gel electrophoresis (SDS-PAGE), 100 ng of each purified protein was mixed with an appropriate volume of 2x Laemmli sample buffer (Bio-Rad) with 5 mM β-mercaptoethanol (Millipore Sigma). Samples were heated at 95 °C for 5 minutes to denature and reduce the proteins and were then separated by electrophoresis through 4-15% Mini-PROTEAN TGX Stain-Free gel (Bio-Rad) with Tris/Glycine/SDS running buffer. Gels were subsequently stained with Oriole fluorescent gel stain (Bio-Rad) and visualized on a UV transilluminator 2000 (Bio-Rad)

For Western blot, the total protein concentration was determined using the bicinchoninic acid (BCA) protein assay kit (Pierce/Thermo Fisher Scientific; Rockford, IL). 25µg of protein samples were mixed with 4x LDS sample buffer (Life Technologies, CA, USA) and 10x reducing agent (Invitrogen; Carlsbad, CA) and heated for 5min at 100°C. Samples were loaded in a 4-12% SDS-PAGE gels (NUPAGE^TM^; Invitrogen) for electrophoretic separation and subsequently electro-transferred to polyvinyldifluoride (PVDF) membrane (Invitrogen, USA). Membranes were blocked with 5 % bovine serum albumin (BSA) in TBST (tris buffered saline with tween 20) for 1h at room temperature and afterward incubated overnight at 4°C in primary antibodies. Membranes were then washed with TBST (3 times for 15 min) and incubated with anti-rabbit-HRP (111-036-045; Jackson ImmunoResearch; 1:5000) secondary antibody. Membranes were then imaged with Bio-Rad ChemiDoc system (Bio-Rad, Hercules, CA) and analyzed using ImageJ software (NIH). Following primary antibodies were used SARS-CoV-2 membrane protein antibody (ProSci; 9165; 1:1000), SARS-CoV-2 envelope protein antibody (ProSci; 9169; 1:1000), SARS-CoV-2 Nucleocapsid protein (RayBiotech; QHD43423; 1:1000) and SARS-CoV-2 Spike protein antibody (Novus biologicals LLC; NB100-56578; 1:1000).

### Protein reconstitution into vesicles

Monodisperse LUVs were extruded using a vesicle extruder. Lipids with composition mimicking that of the endoplasmic reticulum-Golgi intermediate compartment (ERGIC) were dissolved in chloroform with solid concentration of 5 mg/mL. The molar ratio of different lipids (Avanti Polar Lipids, Inc, Alabaster, AL) are 1-palmitoyl-2-oleoyl-glycero-3-phosphocholine (POPC): 1-palmitoyl-2-oleoyl-sn-glycero-3-phosphoethanolamine (POPE): 1-palmitoyl-2-oleoyl-sn-glycero-3-phosphoinositol (POPI): 1-palmitoyl-2-oleoyl-sn-glycero-3-phospho-L-serine (POPS): Cholesterol = 0.45: 0.2 : 0.13: 0.07: 0.151. The chloroform solution was dried in a glass vial with gentle N2 gas stream, then vacuumed overnight at −30 in Hg at room temperature. The dried lipid mixture was hydrated with biological relevant buffer (150 mM NaCl, 20 mM HEPES, pH = 7.2) with 30 s vortex, prior to ten freeze-thaw cycles in dry ice and 37 °C. After the final thawing step, the aqueous solution was passed 11 times through a polycarbonate membrane with 100 nm pores (Nuclepore Track-Etch membrane, Whatman, Chicago, IL). The size and zeta potential of extruded membrane were measured with dynamic and phase analysis light scattering (DLS and PALS) techniques using a 90 Plus PALS light scattering machine (Brookhaven instruments, Holtsville, NY).

To reconstitute the M proteins into LUV, the concentrated stock solution of n-dodecyl-β-D-maltoside (DDM Avanti Polar Lipids, Inc, Alabaster, AL) were added to the 5 mg/mL of freshly extruded LUVs solution to reach a final concentration of 80 mM. M-protein stabilized by Triton X-100 (stock solution: 1 wt.% Triton X-100 per every 2 mg/mL M protein) was next added to the LUVs solution at a mass ratio of M/lipid = 1/100, 1/67 and 1/50 after 30 min of incubation. The solution was allowed another 30 min of incubation before addition of 40 mg of wet BioBeads (Bio-Rad, Hercules, CA) per mL of LUVs solution to slowly remove the detergent over 3 hours. The second and third doses of BioBeads at 40 mg/mL were also added at 3 hours intervals. A final dose of Biobeads was added at a ratio of 380 mg/mL and incubated for another 3 hours. The M-reconstituted LUVs were separated from the solution via centrifugation at 13 krpm for 30 min in a microcentrifuge tube.

### Preparation of supported bilayer samples

To prepare supported lipid bilayer (SBL) on mica for AFM imaging, 75 μL of LUVs solution collected from the bottom of the microcentrifuge tube was added onto freshly cleaved pristine mica and incubated for 20 min to 1 hour for different coverage. The sample was then gently rinsed with 5 mL of buffer. Caution was taken not to expose the SBL sample directly to air. The samples were kept submerged in aqueous buffer until imaging in an AFM fluid cell. Imaging was performed within 1 hour of the sample preparation.

### AFM imaging, force profile collection and analysis

AFM height images were obtained inside a fluid cell in tapping mode using a MSNL cantilever (Bruker, Camarillo, CA). Images were taken in 512×512 line resolution over areas ranging from 250 nm x 250 nm to 1 μm x 1 μm. The effective tip size was calibrated with 10 nm gold nanoparticles (Ted Pella, Redding, CA) using our previously reported protocol2. Membrane structure was probed with nanoindentation to obtain force profiles. The force profiles were collected in force volume mode with a 16 x 16 sampling resolution over the designated area of interest of 500 nm x 500 nm. To probe the continuum elastic property of the membrane and protein patches, an MLCT probe (Bruker, CA) with a larger calibrated tip radius of 23 nm was used. The area of interest was identified from prior tapping mode imaging. The spring constant of the larger tip radius cantilvers was calibrated to be 0.15 N/m using its thermal oscillation spectrum.

AFM image analysis were performed in Mountains SPIP software (Digital Surf, France). Protein particles were identified by setting the height of SBL top surface as the threshold and only identifying out protrusions above the membrane threshold. The software-identified protrusions were then manually examined to exclude noise and false identification of the protein particles. The false identifications by the software can be ruled out as they are 1-2 pixel wide noise peaks which are much smaller than the proteins which are greater than 40 pixels depending on whether they isolated protein dimers or large patches.

### CryoEM data collection and analysis

Cryo-samples were prepared in a Vitribot Mark IV (Thermo Fisher Scientific). For each sample, Quantifoil grid R2/1 was glow-discharged before a 3.5 μL of aliquot was applied and blotted for 4 s at 95% humidity at room temperature. The samples were then plunged into liquid ethane and preserved in liquid nitrogen. The Cryo-EM micrographs were obtained using a Talos Arctica (FEI) operating at 200 kV equipped with a Falcon 4i Direct Electron Detector (Thermo Fisher Scientific). The imaging was performed at 150,000x magnification with a resolution of 0.95 Å/pixel and a defocus range of −1.5 to −2.5 μm. About 150-250 images were collected for each sample using automated data acquisition using EPU software (Thermo Fisher Scientific). Images were motion corrected within EPU. The data set was processed using Relion software. ∼ 500 areas of interests were manually picked to generate the 2D class average images.

### All Atom Molecular Dynamics Simulation

All atom molecular dynamics (MD) simulations were performed using the CHARMM36m force field with the MD package GROMACS, version 2022.3 ^30, 31^. The CHARMM-GUI input generator was used to set up the simulated systems with periodic boundary conditions and supplied the six steps used for equilibration ^32–40^. After equilibration, each system was simulated for 1 microsecond with a timestep of 2 femtoseconds in the NPT ensemble. An in-depth table for the conditions used during equilibration steps can be seen in Supplementary Information. System temperature was maintained at 303.15 K using the Nose-Hoover thermostat ^41, 42^, with the pressure maintained semi-isotropically at 1 bar in the x-y dimensions and separately in the z-dimension using the Parrinello-Rahman barostat ^43, 44^. The coordinates for each simulation were saved once every 50 thousand timesteps, or every 0.1 ns, for a total of 10 thousand frames.

Three different membrane systems were simulated, one for each form of the M protein dimer (“short” vs “long”), and one as a control only consisting of a membrane. For the simulations involving an M protein dimer, the protein was inserted into the membrane with an orientation and depth that matched other studies ^3, 4^. In the case of the short form, only residues 9-204 were considered (PDB: 7vgs), while residues 9-206 were included in the long form structure (PDB: 7vgr) ^3^. The membrane was composed of Chol 15%; DOPC 45%; DOPE 20%; DOPS 7%; POPI 13% in both leaflets, with the solvent consisting of NaCl at a concentration of 0.15 M and TIP3P water. Each simulation, depending on the inclusion of a protein, consisted of a 25 nm x 25 nm membrane in the x-y plane with at least 5 nm of solvent above and below the protruding protein, yielding a total unit cell thickness of ∼ 18 nm in the z dimension.

For runs involving an M protein, the final trajectory was reoriented frame-by-frame such that the protein was centered in the box for all frames. While each trajectory was fitted to eliminate protein translation across the membrane, this was not the case in the direction normal to the membrane and for the rotation of the protein. To analyze the processed trajectory, the python library MDAnalysis was used ^45, 46^. Additionally, all atom simulations for each conformation of the M protein dimer in solvent were performed to compare protein stability with the membrane simulations. These systems involved a box size of approximately 15(16) nm x 15(16) nm x 15(16) nm for the short(long) conformation. The same process and conditions as the membrane simulations were used, with slight deviations in the equilibration steps described in Supplementary Information.

### Coarse-grained simulation

The coarse-grained simulation is performed using HOOMD-blue package ^47^ and BondFlip plugin ^13^. For the membrane, we initially assign a 100a×84a triangular lattice plane and place M proteins in the middle disk area with a radius of 15a. The dynamics of the membrane is modeled through Langevin integrator with *dt* = 0.001*s* in room temperature. In every 100 steps, we apply Monte-Carlo bond-flip to change the local connectivity of the membrane, where each bond has stretching and bending energy with spring constant 20*k*_*B*_*T*/*a*^2^ and bending rigidity 20*k*_*B*_*T*. We note that each simulation is run for 10,000s, and a height constraint (ℎ(*t*) = 0) is applied for the region with radial distance *r* > 35*a* through the simulation time. The simulation is performed on NVIDIA GeForce RTX 3090, and the visualization is performed with OVITO software ^48^.

## Acknowledgements

This research was funded by the University of California Office of the President UC Multicampus Research Programs and Initiatives, grant number M21PR3267 and the National Science Foundation RAPID grant, 2034794.

## Author Contributions

Conceptualization, T.E.K., M.C., A.G., U.M., and R.Z..; methodology, T.E.K., S.A., Y.Z., J.M., M.W., and S.L.; investigation, Y.Z., S.A., M.W., J.M. S.L., U.M and T.E.K.; writing—original draft preparation, Y.Z., S.L, M.C., J.M., U.M., R.Z., A.G. and T.E.K.; writing—review and editing, M.C., U.M., R.Z., A.G. and T.E.K.; supervision, T.E.K., M.C., A.G., U.M., and R.Z.; project administration, T.E.K., M.C., A.G., U.M., and R.Z.; funding acquisition, T.E.K., M.C., A.G., U.M., and R.Z. All authors have read and agreed to the published version of the manuscript.

## Data Availability Statement

The data presented in this study are available on request from the corresponding author.

