## Supplementary Information for "Synthesis, Insertion, and Characterization of SARS-CoV-2 Membrane Protein Within Lipid Bilayers"

#### SUPPLEMENTARY METHODS

**Determination of detergent conditions for M protein insertion**

The proper function of transmembrane M protein requires insertion into the bilayer membrane with the correct orientation. In this study, the protein insertion utilized a detergent assisted procedure and was optimized for M protein in terms of the type and concentration of detergent used. Fig. S1 shows the key results obtained during the optimization process. The as-extruded lipid vesicles were measured to exhibit an intensity-averaged hydrodynamic size of 123 nm in DLS (Fig. S1a). As shown in Fig. S1b, as detergents are gradually introduced into the vesicle solution, the lipid vesicles undergo a three-stage evolution, which has been well established in previous studies [1,2]. In the first stage, the detergent molecules insert themselves into the lipid vesicle bilayer structures without dissolving the vesicles. In this stage, with more detergent addition, the detergent concentration in the bilayer membrane increases gradually without decreasing the size and number of vesicles. At the saturation concentration, $C_{sat}$, the vesicle becomes saturated with detergent. Increasing the detergent concentration beyond saturation leads to the second stage where further detergent addition results in the partial dissociation of the original vesicles. Here, the hydrodynamic size of vesicles significantly decreases with detergent addition, as they are being depleted of lipids. The dissociated lipid molecules form mixed micelles with the detergents. At the end of the second stage, all the vesicles are dissociated completely at the vesicle dissolution concentration, $C_{sol}$. In this third stage, only mixed micelles exist. Further addition of detergents only increases the detergent composition in the mixed micelles.

In this study, two non-ionic detergents, TX-100 and DDM were tried. Unlike their ionic counterparts, these detergents are mild and do not disrupt protein folding [1]. Figs. S1c,d show the hydrodynamic size change with detergent concentration for TX-100 and DDM, where the three stages described in Fig. S1b were observed for most cases. TX-100 was found to saturate and dissolve vesicles at much lower detergent concentrations compared with DDM. Fig. S1e,f show the saturation and dissolution concentrations as a function of the total lipid concentration in solution for TX-100 and DDM. In both cases, saturation and dissolution concentrations were found to increase with increased lipid concentration. For the concentrations studied TX-100 shows a very gradual linear increase while DDM the saturation concentration increases rapidly and then reaches a plateau at higher lipid concentrations. Thus, DDM provides much better control and higher tolerance and therefore for the M protein insertion into the lipid bilayers. The DDM concentration used for M protein insertion was ~80% of $\boldsymbol{C_{sat}}$. A similar detergent (octylglucoside) was reported to demonstrate optimum transmembrane protein function and unidirectional insertion in this concentration regime [3]. The unidirectional insertion has been verified by the CryoEM and AFM results as reported below.

After M protein incubation, the detergent removal rate was also optimized. The detergent was removed via hydrophobic adsorption, where hydrophobic polystyrene resin [4] (SM-2) bio beads at a 40 mg/mL were added into the solution. This lower bead concentration in contrast to the usual 80 mg/mL lead to a slower detergent removal rate, which facilitated a higher efficiency in the protein insertion.

**Determination of elastic moduli from AFM measurements**

Fig. S2 shows the average elastic moduli extracted from nanoindentation force profiles for SLB and the M protein patches. SLB exhibits a Young's modulus of 9.5 MPa, which agrees with literature values [5]. The M protein patches demonstrated a much higher modulus value of 41.5 MPa, indicating the patches are indeed a different type of material from the lipid bilayer.

AFM nanoindentation was performed in force volume mode using a MLCT probe (Bruker, CA). The spring constant was calibrated using a thermal tune, $k$ = 0.15 N/m. Tip radius was calibrated with 10 nm gold colloids (Ted Pella, CA), $R$ = 23 nm. The force profiles were collected over a 1 μm x 1 μm area at a 16 x 16 resolution. To extract the Young's modulus of SLB and the protein patches, a linearized Hertzian contact model was utilized. According to Hertzian model, the relationship between measured force and sample indentation can be expressed as follow:

$$F = \frac{4ER^{1/2}}{3(1-\sigma^2)}\delta^{3/2}.$$

Here, $F$ is the force measured from AFM (Hook's law, $F$ = cantilever deflection x calibrated spring constant). $R$ is the calibrated tip radius of AFM probe, σ is the Poisson's ratio. Here, both $\boldsymbol{\sigma_{SLB}}$ and $\boldsymbol{\sigma_{Mprotein}}$ are taken to be 0.5, as most biological materials such as lipids and proteins are considered incompressible. $\delta$ is the indentation depth. The acquisition of indentation depth is relatively complicated considering it requires the identification of exact point of tip-sample contact. However, the tip-sample separation, $D$, can be easily obtained as raw data and the relationship between $D$ and $\delta$ is simply:

$$D = c - \delta.$$

Here, $c$ is a constant. To simplify the analysis, the Hertzian equation was linearized [6] as follows:

$$F^{2/3} = C - D * [\frac{4}{3}\frac{ER^{1/2}}{(1-\sigma^2)}]^{2/3}.$$

Here, $C$ is another constant. Therefore, by obtaining the slope of the linear relationship between $F^{2/3}$ and $D$, we can extract the elastic modulus. Here, to rule out the substrate effect and make sure the $E$ is extracted from the linear elastic region, only the initial ~ 20% of force curve was used to be analyzed by the Hertzian model.

**Atomistic Molecular Dynamics (MD) simulations of M-protein stability:**

Two interconvertible forms of the M protein homodimer have been reported for SARS-CoV and SARS-CoV-2 [7,8]. To investigate the relative stability of these forms we performed MD simulations of both the long and short forms based on Cryo-EM structures [7]. We ran these simulations in two different environments, embedded within a membrane bilayer (as shown in Fig. S3) and in an environment of just water and counter ions. The latter "solvent" simulations will help determine if the relative stability of the long and short M protein dimer conformations is a result of interactions with the membrane or are intrinsic to the structures themselves. While other M protein dimer simulations using experimentally determined structures have found these conformations to be stable in the membrane, the periodic box size for those simulations was either very small or the long form was not considered [7,9].

From these simulations, the long form in solvent changes most drastically from its initial configuration. The changes in the structural conformation of the short form in solvent and membrane are within the level typical of thermal fluctuations (~0.35 nm) as shown in Fig. S4a. In contrast, the solvent-phase long form RMSD is nearly double that observed for the short form solvent-phase simulation after 1 microsecond of simulation (0.7 vs 0.35 nm). In the membrane, the long form structure has a final RMSD that is approximately 30% larger than the short form in membrane. We see a similar result for the radius of gyration ($R_g$), where the long form in solvent's $R_g$ decreases over the course of the simulation to match the $R_g$ observed for the short form in both the membrane and solvent. The $R_g$ also decreases in the long form membrane simulation, however, it does not drop to a level comparable to the solvent simulation.

These results suggest that the long form in solvent is not at a minimum energy configuration initially, unlike the short form, which undergoes little change throughout the simulation. The N-terminal region of the protein curls up without the membrane present to prevent it for the long form. On the other hand, when embedded in the membrane, the alpha helices were split apart by neighboring lipids. Of particular importance is the W31 residue mentioned in Zhang *et al.* [7], which rotates away from its duplicate due to interactions with nearby DOPC. In comparison to the simulations Zhang et al. ran, these were not performed with the WYF parameter [10]. However, since the nearby DOPC lipids interact with W31 throughout the simulation, including the WYF parameter would only increase this interaction and would not prevent the alpha helices from splitting apart [11]. This supports the suggestion that the long form of the M protein dimer needs both the membrane and possibly other proteins to stabilize its structure [8].

**MD simulations of M protein effects on membrane thickness:**

According to the MD simulations, the short and long conformations alter the thickness of the membrane adjacent to the protein differently. Average values of the thickness, calculated as the difference in height between the phosphate groups in the lower and upper leaflets over the last 500 ns of the simulation, can be seen in Fig. S6. At each point in time every phospholipid head was placed into a bin (the membrane was split into 15 x 15 bins), where the average upper leaflet height and lower leaflet height for each bin were subtracted from each other. This process was performed using MDAnalysis.

Based on the line of best fit and the corresponding 95% confidence intervals, the long and short form induce statistically significant changes in membrane thickness that depend on the radial distance from the protein. In contrast, the membrane-only simulation thickness appears to be independent of the distance from the center, hovering around ~4.20 nm. Thus, the membrane-only system experiences thickness fluctuations, while simulations with M protein exhibit persistent thickness dependencies radially around the embedded protein dimer.

Combining these results with the radially binned averages represented with the black line of each bottom plot, the long form of the protein is surrounded by a region of thicker membrane, while the short form is surrounded by a region of thinner membrane. This agrees with Neuman *et al.* [8], and their study of M proteins during the formation of viral particles, in which they found long form in thicker regions. These changes are possibly induced by the different shape and/or different membrane-facing surface areas between the two conformations. Alternatively, the different M protein forms may simply stabilize different membrane thicknesses arising due to membrane fluctuations.

**Impact of M protein on membrane curvature in MD simulations**

To decouple the height of the membrane from the thickness, membrane midplane height is analyzed over the last 500 ns of the simulation. The midplane is defined as the average surface between the lower and upper leaflet shown in Fig. S3c. As seen in Fig. S7, while the membrane-only simulation has slight height variation, it is clearly not as substantial as either the short or long form runs. Thus, it can be assumed that these represent membrane fluctuations that persist on the scale of 500 ns.

From the line of best fit, with the given 95% confidence intervals for each simulation, the long conformation of the M protein dimer seems to have the greatest effect on membrane deformation. Additionally, since the confidence interval does not include the horizontal (slope = 0), this radial dependence of average membrane midplane height appears significant.

The shape of the membrane curvature can be determined from the radial average. While the deformation of the membrane-only simulation matches that of height fluctuations, both membranes containing either the long or short M protein conformations seem to adopt persistent curvature, in the form of a valley-like fold in the membrane enclosing the protein dimer. This shows that the M protein induces curvature in the membrane. Due to the small size of the lipid membrane patch simulated and the periodic boundary conditions imposed, this is only a qualitative confirmation of the membrane curvature induced by the protein. As discussed in Neuman *et al.* [8] for SARS-CoV, the long conformation appears to induce more curvature than the short conformation, agreeing with our observed membrane deformation.

**Protein rotation in MD Simulations**

As seen in the superposition of protein cross sections throughout the simulation in Fig. S6 and S7, the short M protein dimer conformation appears to have a more circular cross section than the long form. Our analysis shows this is a result of the short form’s increased rotation in comparison to the long form. This rotation can be measured through the calculation of the protein's inertia tensor, and subsequent computation of the dot product between the x-axis and an inertia eigenvector. Performing this analysis on both the short and long form leads to the results shown in Fig. S8.

As seen in Fig. S8b-c, the short conformation rotates over a wider angular range than the long conformation. The range of rotation is twice as large for the short form than the long form with several peaks in the frequency plot. For the case of the long form, the protein is aligned with the trough of the membrane, which could be an explanation for the bend's formation: that different axes for the protein induce different levels of curvature. Furthermore, the bending aligned with respect to the long axis of the protein can imply a possible process for the formation of higher order oligomers as other M protein dimers align in this trough. Imaging studies of the M protein for SARS-CoV have shown line-like formations across the virion [8,12].

**Coarse-grained Molecular Dynamics Simulation of M protein effects on membrane curvature**

The role of M Proteins in inducing membrane curvature and budding in ERGIC was previously explored [13]. ERGIC was modeled as a sphere built from a triangular lattice as shown in Fig. S12, which shows how the M-M interaction can result in the budding of the region of the ERGIC membrane where the M proteins are embedded. The ERGIC is shown as a gray vesicle in Fig. S12 and the three regions of M protein are depicted with three hard spheres (M1 in red, M2 in orange, and M3 in black), where M1 and M2 correspond to the transmembrane domain, and M3 denotes the endodomain. Figure S12a shows that in the absence of endodomain interaction, the membrane does not curve. However, the attractive interaction between endodomains results in the membrane curvature and budding as illustrated in Fig. S12b. However, in the absence of interaction between endodomains, the attractive interaction between N proteins (blue and yellow beads) and M proteins, see Fig. S12c, or between RNA and M proteins (Fig. S12d) can induce curvature and budding.

Since most of the experiments and all-atom simulations in this paper are performed on the flat membrane, for the purpose of this paper, we simulated the role of M proteins embedded in a flat membrane, see the main text. Previously, it was found that the curvature of ERGIC helps with overcoming the energy barriers and leads to budding. In this paper, we found that M proteins can curve even the flat membranes and induce budding if the endodomains interact with each other.

**Estimation of line tension from membrane thinning**

For a given height mismatch, δ, between a domain and the surrounding lipid membrane of thickness h, the line tension generated due to the elastic cost of bend and tilt deformation is given by

$$\gamma=(\delta/h)^2\sqrt{B_sK_s}\sqrt{B_rK_r}\,/(\sqrt{B_sK_s}+\sqrt{B_rK_r}),$$

where $B_{s,r}$ and $K_{s,r}$ are the bending and tilt modulus of the soft (lipid)/ rigid (domain, M protein) phases respectively. We can use this expression for a monolayer assuming the height mismatch is evenly distributed among the two leaflets. Since the stiffness of the M protein region (41.5 MPa) is much higher than that of the lipid layer (9.5 MPa), we can take $B_r >> B_s$ and $K_r >> K_s$, simplifying our expression to

$$\gamma=\left(\frac{\delta}{h}\right)^2\sqrt{B_SK_S}$$

We can estimate the bending modulus $B_s$ from the measured Young's modulus via

$$B_s = Yh^3/24(1-\nu^2),$$

With $\nu$ = 0.5 (incompressible) as approximately 8 $k_BT$ and we can take $K_s$= 8 $k_BT/\mathrm{nm}^2$, which is consistent with values derived from simulations and experiments [14]. Using these moduli along with δ = 0.5 nm and h = 4 nm, in the simplified expression for the line tension gives $\gamma$ = 0.5 pN.

### SUPPLEMENTARY FIGURES

### FIGURE S1

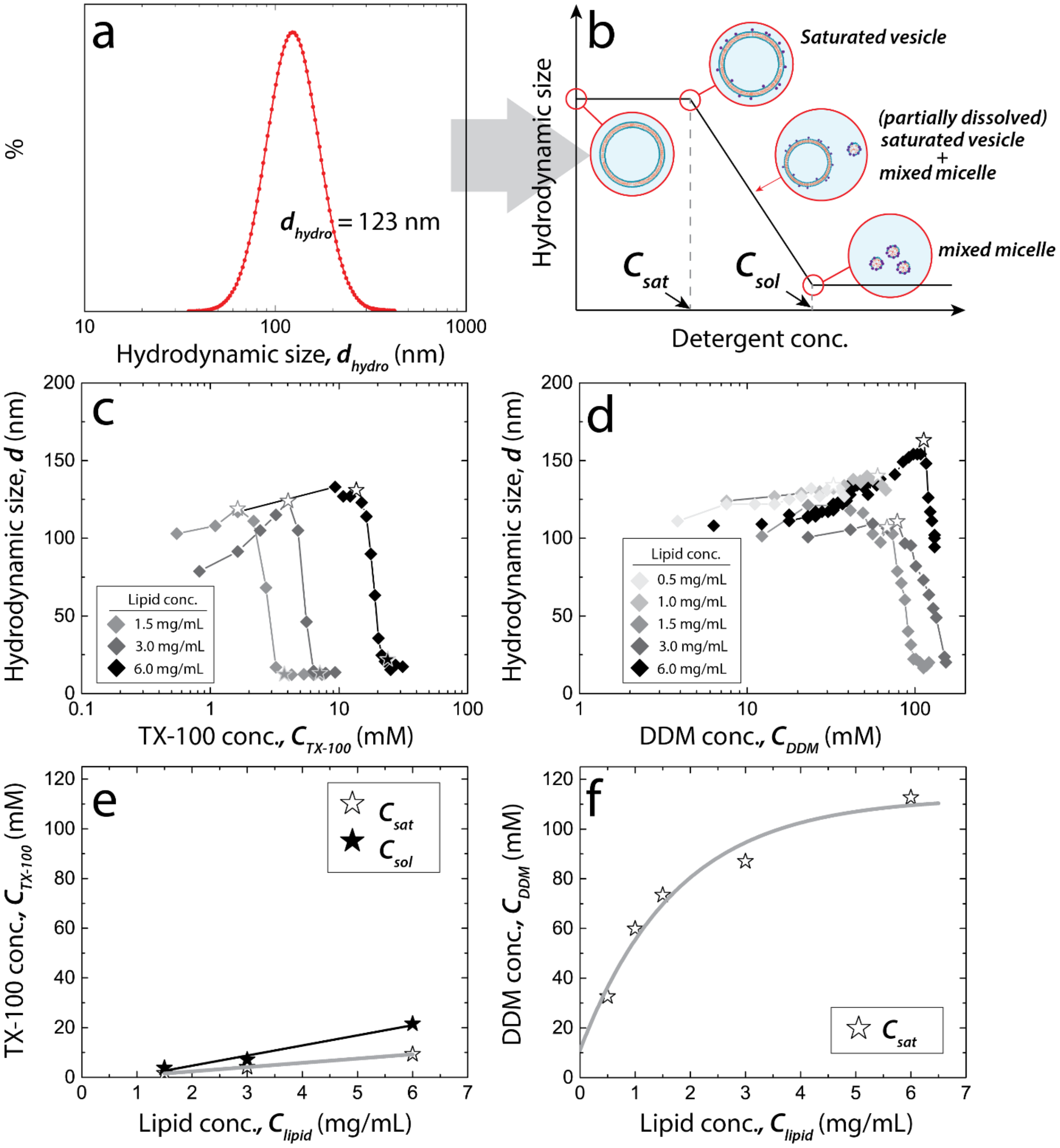


**Figure S1.** Dynamic light scattering (DLS) study of vesicle size as a function of detergent concentration. (**a**) Intensity distribution vs. hydrodynamic size measured with DLS for as-extruded lipid vesicles in biological buffer. (**b**) Schematics of the three stages of vesicle dissolution with increasing detergent concentration. (**c**) and (**d**) are hydrodynamic size of vesicles measured as functions of TX-100 and DDM detergent concentrations, respectively. The empty and filled stars represent the detergent-saturation concentration, $C_{sat}$ and vesicle dissolution concentration, $C_{sol}$. (**e**) and (**f**) are $C_{sat}$ and $C_{sol}$ observed at different lipid concentrations for TX-100 and DDM, respectively.

**FIGURE S2**

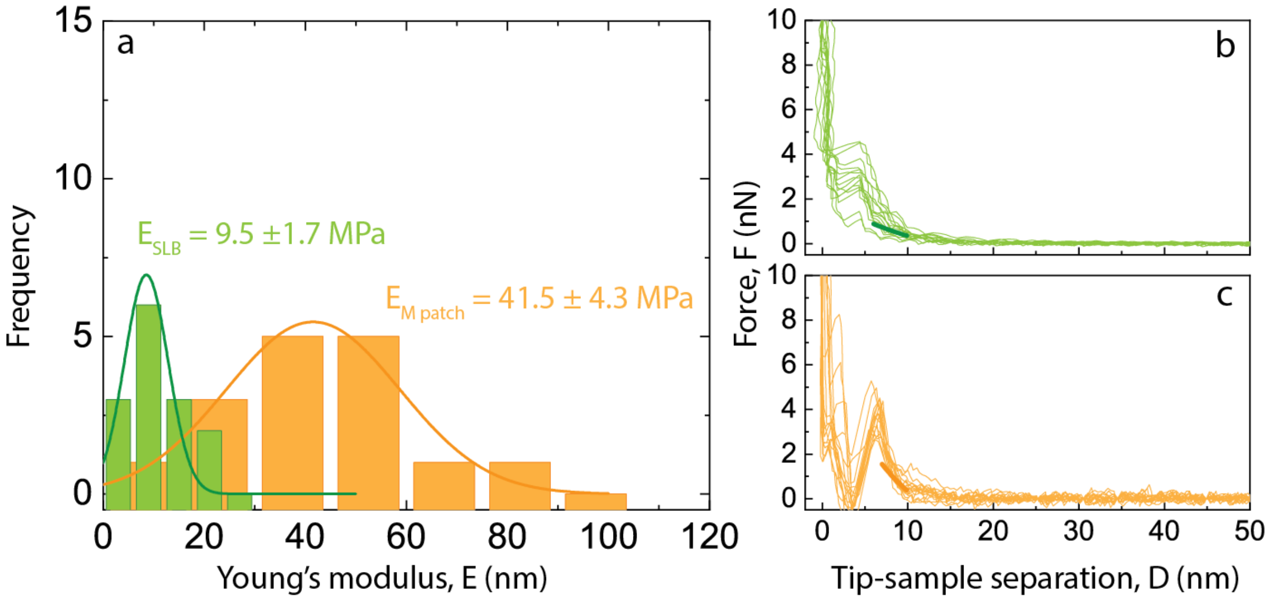


**Figure S2. Elastic modulus of SLB and M protein patches obtained from nanoindentation technique.** (**a**) Histogram of Young's modulus obtained from analyzing the AFM nanoindentation force profiles using Hertzian contact model. (**b**) and (**c**) are the force profiles obtained from SLB and protein patches, respectively. The thick solid lines represent the force profile calculated using the mean elastic modulus values from (a).

**FIGURE S3**

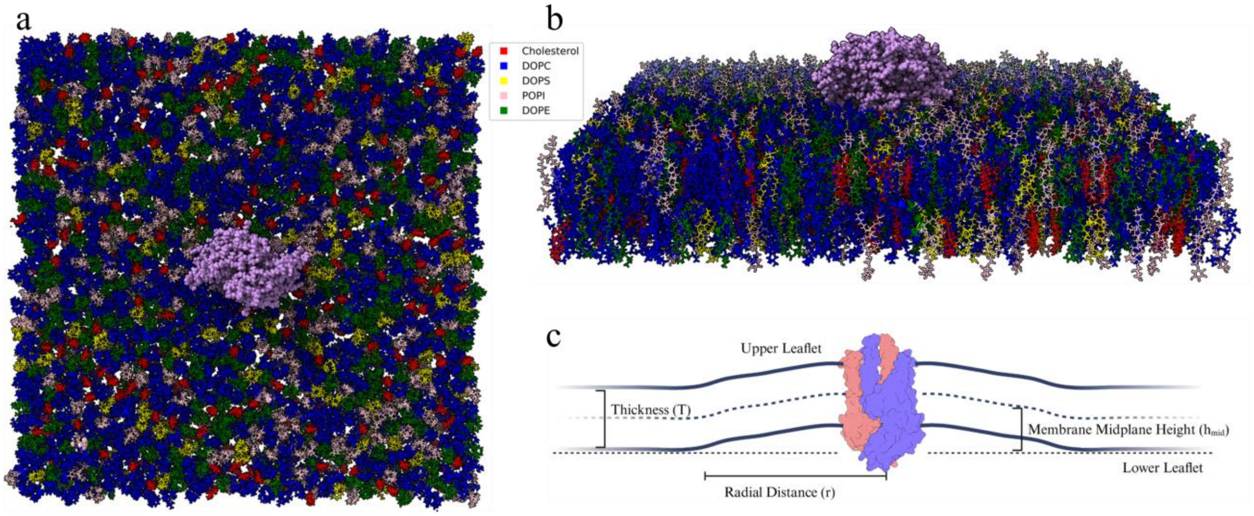


**Figure S3: Initial simulation schematic for the short form embedded in a multicomponent ~ 25 nm x 25 nm membrane.** (**a**) A view from above looking down on the lower leaflet, with the elliptical shape of the C-terminal shown in purple. (**b**) A view from the side. (**c**) Diagram of a long form M protein dimer embedded in a membrane, where the differing protein colors refers to the monomers within the dimer (created with BioRender.com).

**FIGURE S4**

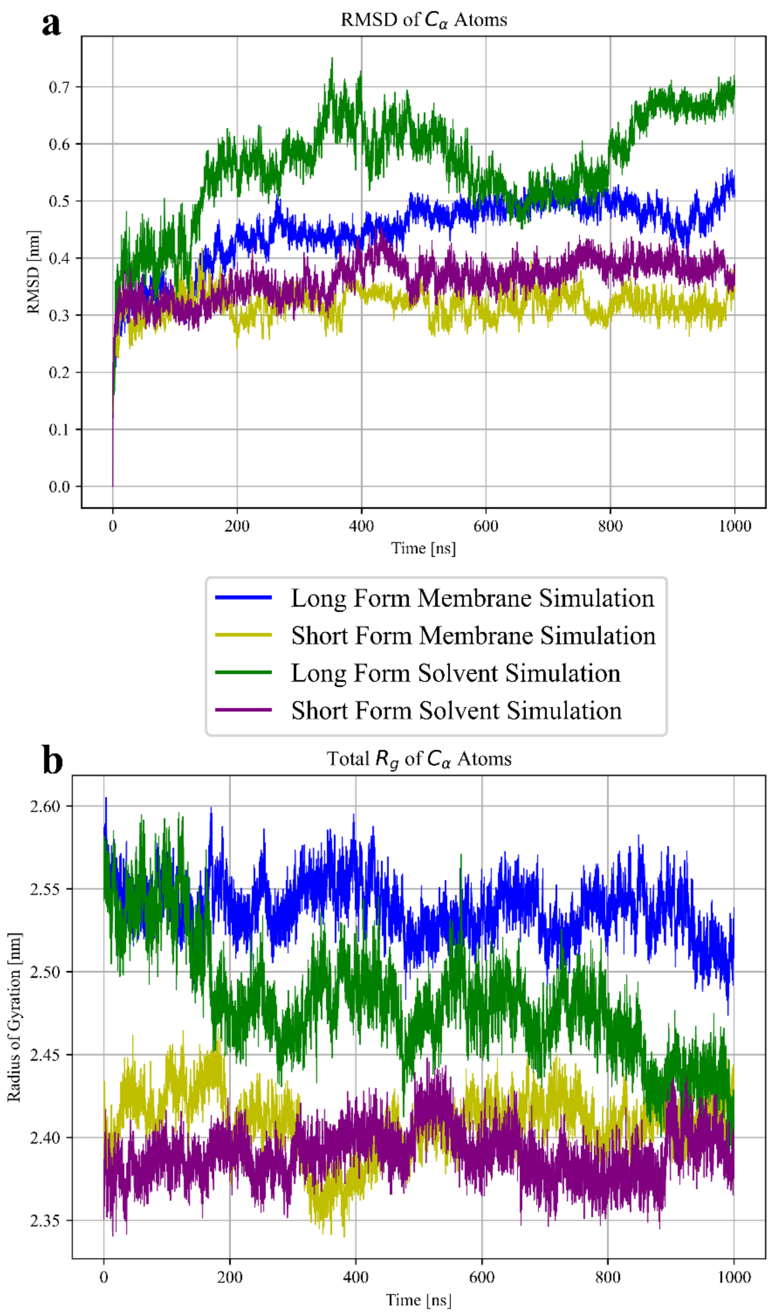


**Figure S4: RMSD and $R_g$ of four different M protein simulations**. The long/short (blue/yellow) form membrane simulations and long/short (green/purple) form solvent simulations. (**a**) Root mean square distance (RMSD) represents the difference of each $C_\alpha$ atom at the corresponding time with its original position squared, summed over every $C_\alpha$ atom, and then divided by the number of $C_\alpha$ atoms and taken to the half power. (**b**) Total radius of gyration $R_g$, defined as the radial distance to a point in which the moment of inertia would match the actual inertia if the body was condensed there, representing how compact the protein is.

### FIGURE S5

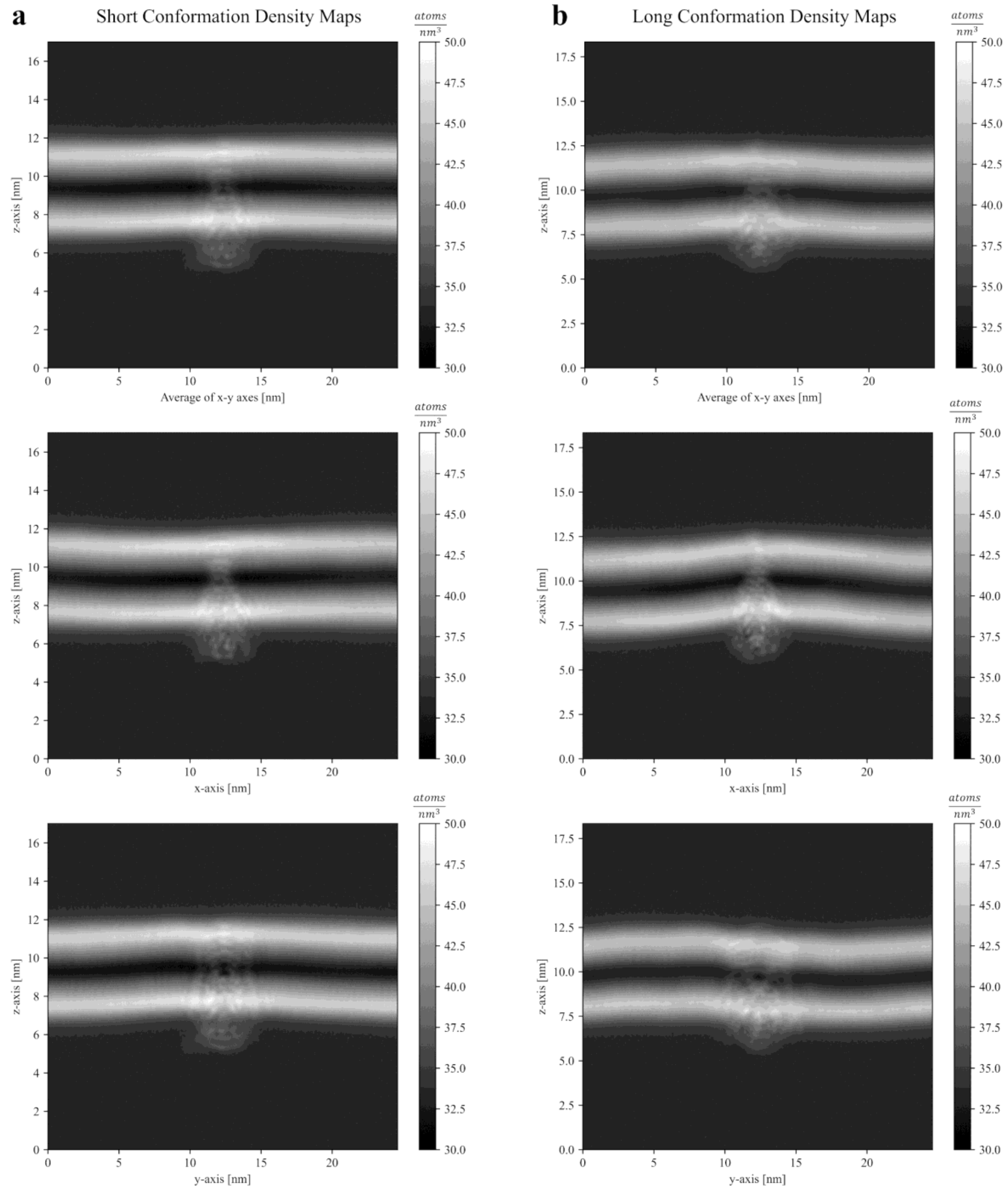


**Figure S5: Atom number density, excluding hydrogen atoms, from MD simulations for the short and long forms**. Generated using the gmx densmap command from GROMACS, (**a**) and (**b**) represent atom number densities for the short and long forms respectively. The whole simulation temporally and spatially is considered, where every atom other than hydrogen is accounted for, with a bin size of 0.095 nm. Each row represents a different viewing angle, with the top an average of the below two.

### FIGURE S6

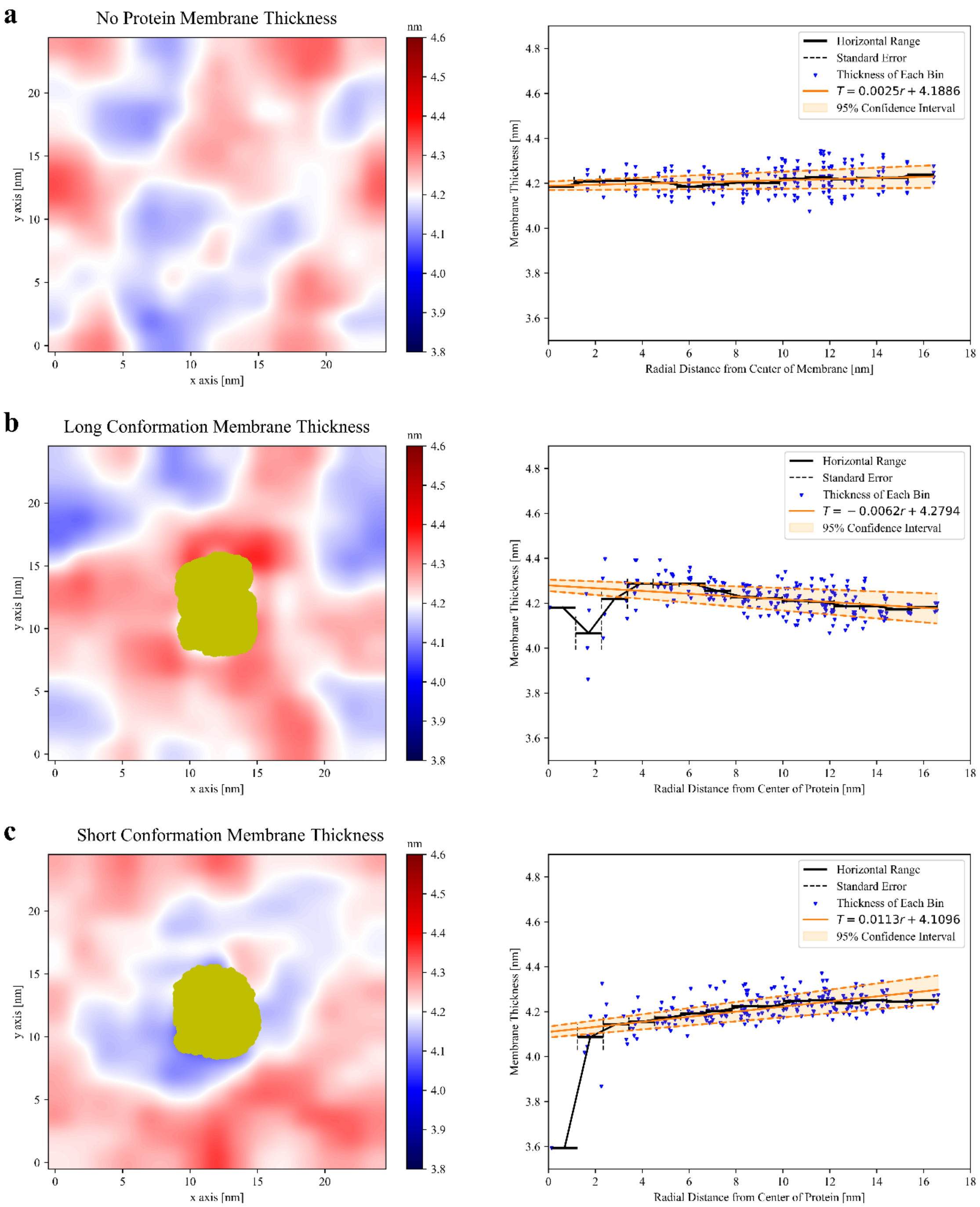


**Figure S6: The average membrane thickness is shown for the last 500 ns of each simulation.** (**a**) – (**c**) Visual from above in nm, with a radial plot from the center of the box or center of the protein shown to the right. Each blue dot represents the average thickness at a given bin, orange defines the 95% confidence interval for a linear line of best fit, and black is a radial average of average thickness. Every atom in the protein at every frame averaged over is shown in gold. (**c**) Shows the non-shifted radial plot used in Fig. 6e.

**FIGURE S7**

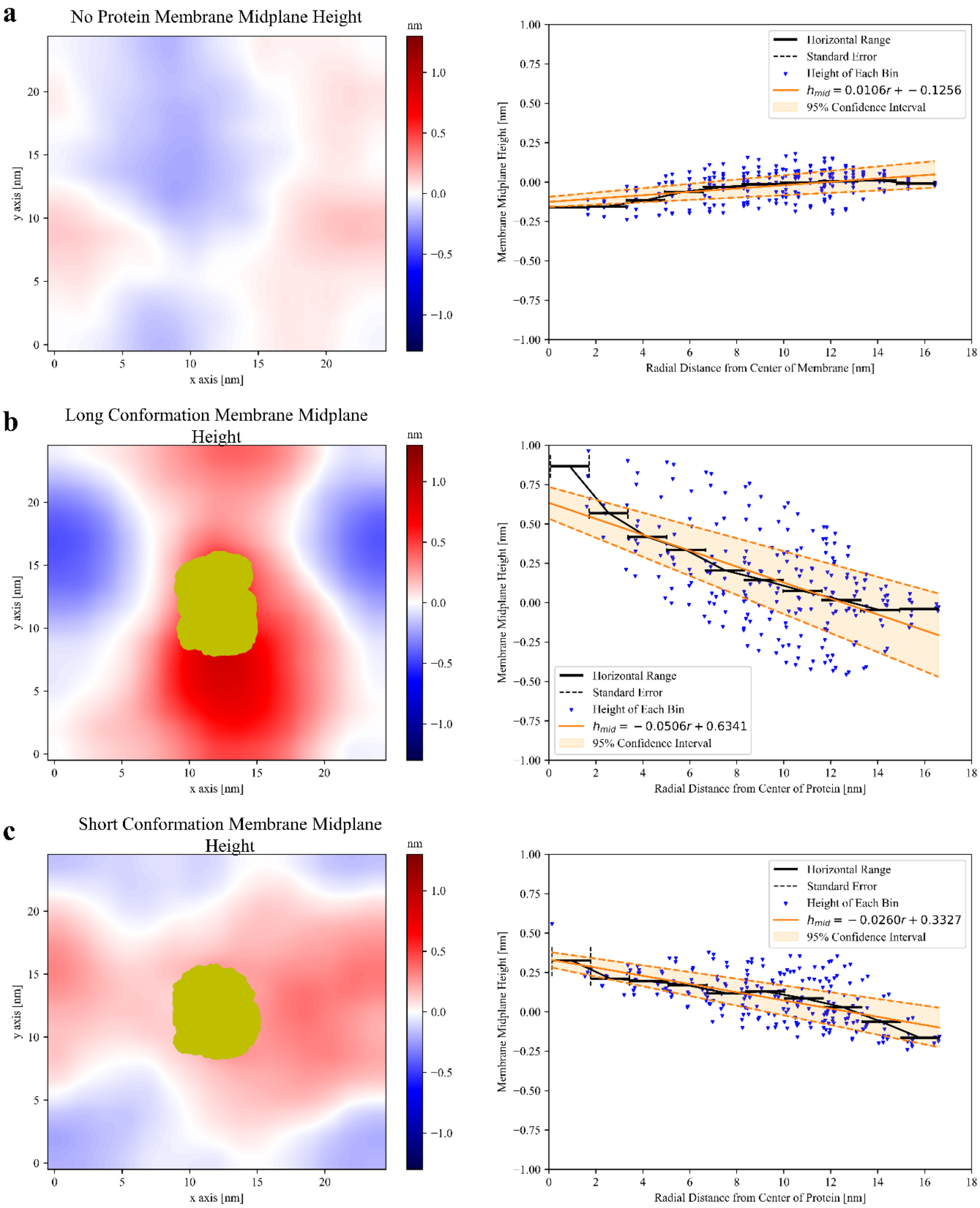


**Figure S7: The average membrane midplane height is shown with respect to the average value along the boundary.** (**a**) – (**c**) Visual from above in nm, with radial plots to the right showing the average heights of each bin (blue) in comparison to either the radial distance from the center of the membrane or the radial distance from the center of the protein. Every atom in the protein at every frame averaged over is shown in gold.

**FIGURE S8**

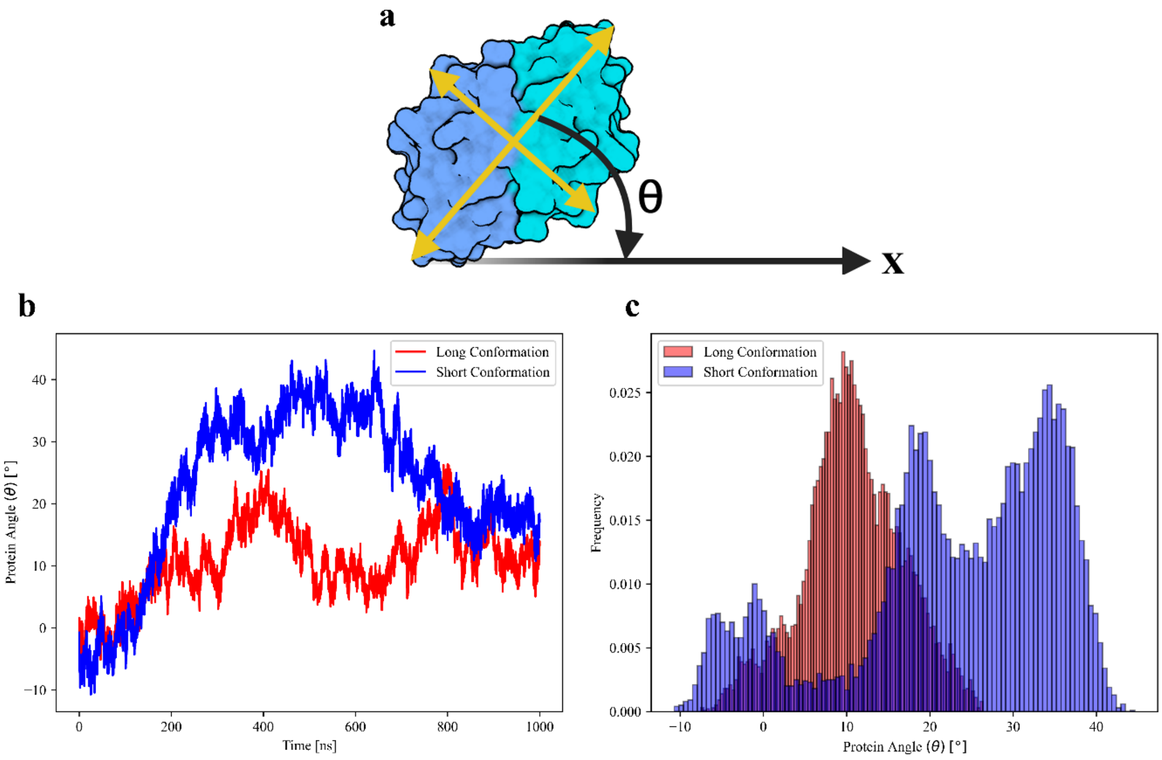


**Figure S8:** Protein rotation for both forms relative to principal axis of the corresponding M protein throughout MD simulations. (**a**) Diagram displaying rotation in the x-y plane, where $\theta$ defines the angle between a principal axis of the protein with respect to the x-axis. The yellow lines represent projections of the long and short axis of the protein. (**b**) Time evolution of the long/short (red/blue) form orientation (θ) with respect to the x-axis. (**c**) Shows the frequency of angles throughout the simulation for both short and long forms.

**FIGURE S9**

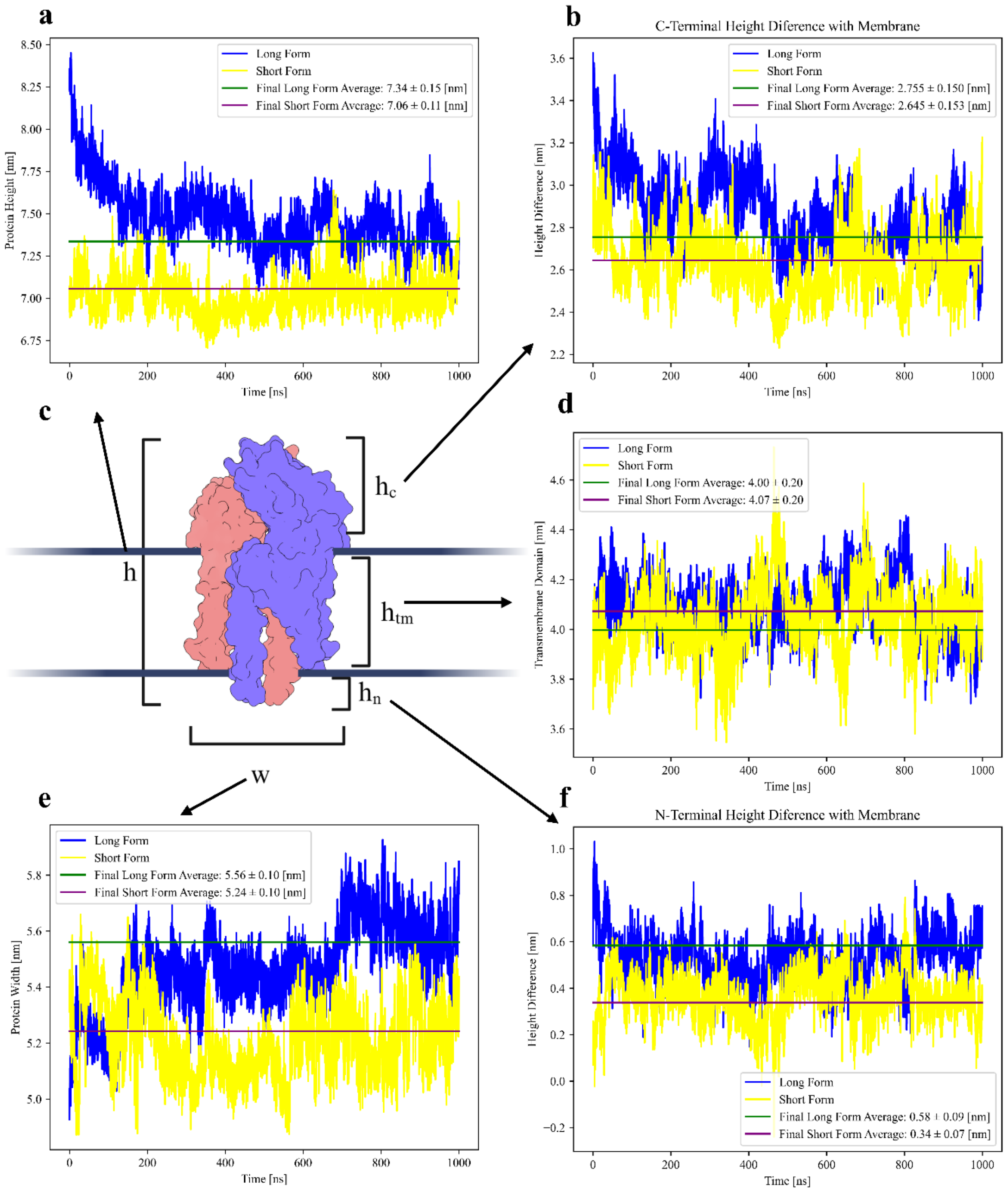


**Figure S9: Long and short conformation dimensions obtained throughout MD simulations.** These calculations were performed with MDAnalysis. (**a**) Protein height, determined by finding the difference between an average of the five highest and lowest $C_\alpha$ atoms throughout the simulation. (**b**) The difference in height between an average of the five furthest away C-terminal $C_\alpha$ atoms and the phospholipid heads within 2 nm of the protein. (**c**) Schematic showing the different characteristic quantities. (**d**) Transmembrane domain height, determined by adding (b) and (f) and subtracting (a). (**e**) Width of the protein, the maximum distance between $C_\alpha$ atoms of like residues. (**f**) The height difference between the N-terminal and the membrane, calculated similarly to (b). Averages were taken over last 100 ns, where the corresponding error is standard deviation.

**FIGURE S10**

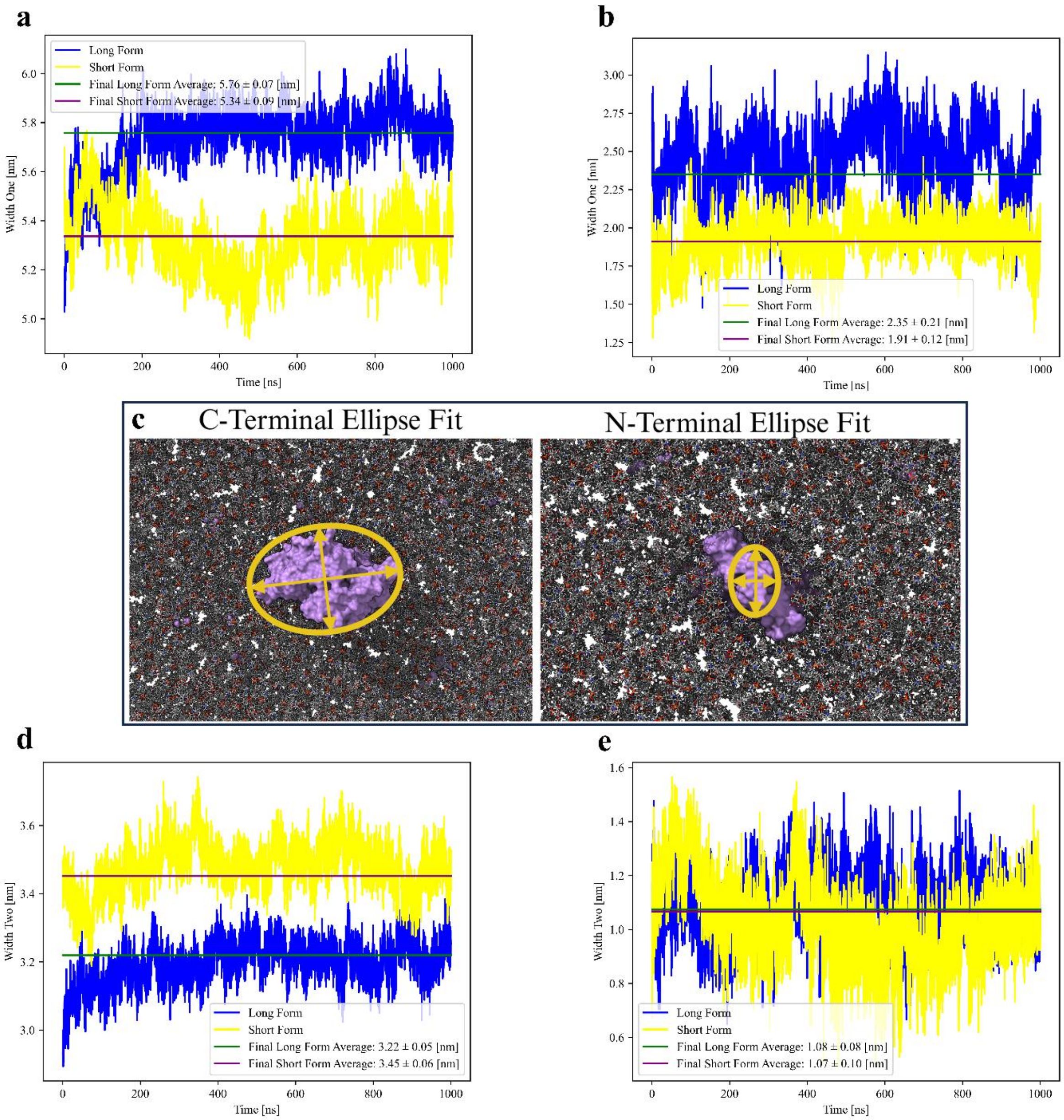


**Figure S10: Dimensions of the M protein when fitted to an ellipse.** (**a**)-(**b**) and (**d**)-(**e**) were obtained by calculating the inertia tensor of every protein atom above the height of the corresponding leaflet, diagonalizing it, and then assuming a uniform protein atom density in the shape of an ellipse. (a)-(b) correspond to the long axis of the C-terminal and N-terminal respectively, while (d)-(e) is the short axis of the C-terminal and N-terminal. (**c**) A schematic of the principal axes of the protein, with the ellipse fit for each terminal shown for the long form of the M protein. Averages were taken over last 100 ns, where the corresponding error is standard deviation.

**FIGURE S11**

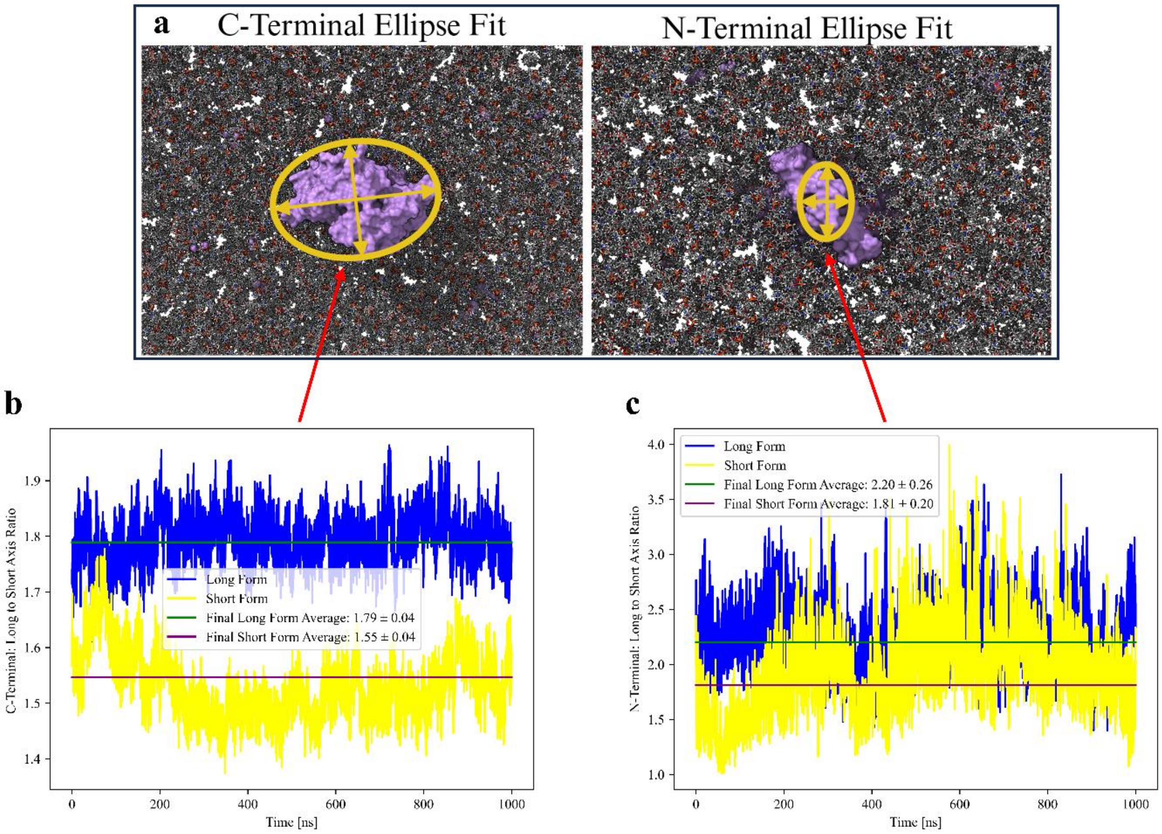


**Figure S11: Long to short axis ratios from M protein ellipse fit.** (**a**) Repeating the schematic shown in Fig. S10. (**b**) Ratios of the C-terminal fit, Fig. S10a and S10d, and (**c**) ratio of the N-terminal fit, Fig. S10b and S10e. Averages were taken over last 100 ns, where the corresponding error is standard deviation.

**FIGURE S12**

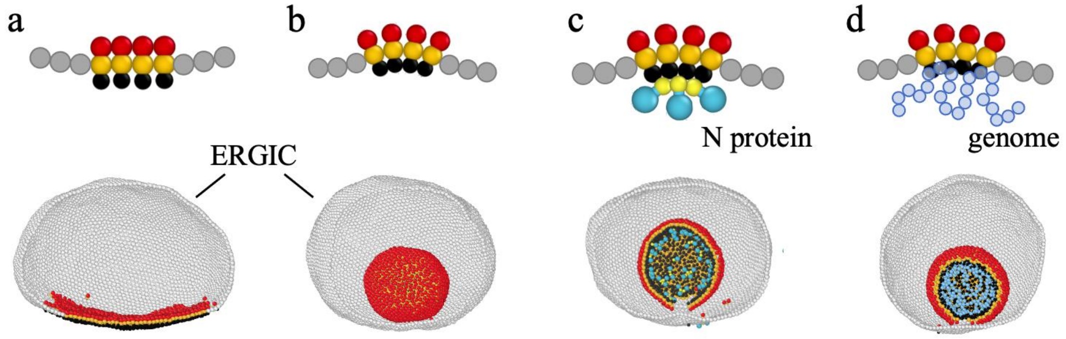


**Figure S12.** Coarse-grained simulations of M proteins embedded in the ER-Golgi intermediate compartment (ERGIC). (**a**) M proteins embedded in the bottom of ERGIC have no endodomain interactions, and no budding is observed in this scenario. (**b**) M proteins have endodomain interactions, proteins bud into ERGIC and form a spherical shell in 4000s. (**c**) M proteins have no endodomain interactions, a budding process happens when N proteins are added. Note that N proteins interact with M proteins endodomain and effectively introduce endodomain interactions. (**d**) M proteins have no endodomain interactions, a budding happens when a short RNA genome (Length = $400a$) is added. The M proteins interact with the genome through screened electrostatic interactions, which effectively cause endodomain interactions.
